# AAV-delivered gene editing for latent genital or orofacial herpes simplex virus infection reduces ganglionic viral load and minimizes subsequent viral shedding in mice

**DOI:** 10.1101/2022.09.23.509057

**Authors:** Martine Aubert, Anoria K. Haick, Daniel E. Strongin, Lindsay M. Klouser, Michelle A. Loprieno, Laurence Stensland, Tracy K. Santo, Meei-Li Huang, Ollivier Hyrien, Daniel Stone, Keith R. Jerome

## Abstract

Herpes simplex virus (HSV) establishes latency in ganglionic neurons of the peripheral nervous system, from which it can reactivate, causing recurrent disease and possible transmission to a new host. Current anti-HSV therapy does not eliminate latent HSV, and thus is only suppressive rather than curative. We developed a potentially curative approach to latent HSV infection and pathogenesis, based on gene editing using HSV-specific meganucleases delivered by adeno-associated virus (AAV) vectors. Our results demonstrated that a dual meganuclease therapy, composed of two anti-HSV-1 meganucleases delivered by a triple AAV serotype combination (AAV9, AAV-Dj/8, AAV-Rh10), can eliminate up to 97% of latent HSV DNA from ganglia in both ocular and vaginal mouse models of latent HSV infection. Using a novel pharmacological approach to reactivate latent HSV-1 in mice with the bromodomain inhibitor JQ-1, we demonstrated that this reduction in ganglionic viral load leads to a significant reduction of viral shedding from treated *vs*. control mice, with many treated mice showing no detectable virus shedding. In general, therapy was well tolerated, although dose-ranging studies showed hepatotoxicity at high AAV doses, consistent with previous observations in animals and humans. Also in agreement with previous literature, we observed subtle histological evidence of neuronal injury in some experimental mice, although none of the mice demonstrated observable neurological signs or deficits. These results reinforce the curative potential of gene editing for latent orofacial and genital HSV disease, and provide a framework for additional safety studies before human trials can begin.

## INTRODUCTION

HSV-1 and HSV-2 were introduced into human ancestors roughly 6 and 1.6 million years ago respectively (Wertheim et al. 2014), and infect a substantial fraction of the global population (about 66.6 % for HSV-1, and 13.2 % for HSV-2) (James et al. 2020). HSV infections can cause recurrent oral, anogenital, or other lesions, while infections of newborns can lead to disseminated disease and devastating neurological sequelae.

Genital infection with HSV-2 increases the risk of acquisition of HIV, and as such genital infection is a major driver of the global HIV pandemic (Schiffer and Corey 2009). While current antiviral therapies can treat acute episodes and suppress outbreaks, they do not cure established infection (Corey et al. 2004; Gupta et al. 2004; Fife et al. 2006; Mertz et al. 1988; Wald et al. 1996).

The characteristic recurrent nature of HSV infection results from the ability of HSV to establish long-term latent infection within ganglionic neurons that innervate the affected sites. This latent HSV in ganglia is unaffected by traditional antivirals, explaining the inability of antivirals to cure, and reactivations typically commence again once therapy is stopped. One of the most promising approaches to a potentially curative therapy for HSV involves gene editing directed at the latent forms of the virus itself. In a recent study, AAV-delivered meganucleases were able to eliminate over 90% of HSV-1 from the superior cervical ganglia of latently infected mice (Aubert et al. 2020).

Despite the impressive reduction of ganglionic HSV loads seen after gene editing in mouse model of orofacial infection (Aubert et al. 2020), the relevance that such reduction would have for human HSV infection is uncertain. Infected persons are typically not concerned with their ganglionic viral loads *per se*, but instead are most concerned with symptomatic disease and/or viral shedding, with its associated risk of transmission to others (Oseso et al. 2016). Unfortunately, existing mouse models are limited in their ability to address these issues; while mice establish latent HSV infection, they rarely spontaneously reactivate virus or show viral shedding at peripheral tissues. Here we introduce a new model of small-molecule induction of HSV-1 reactivation and peripheral shedding in latently infected mice, and use it to demonstrate that gene editing can mediate a dramatic reduction not only in ganglionic viral loads, but also in induced orofacial and genital viral shedding after AAV-delivered meganuclease therapy.

## METHODS

### Mice

Six-to eight-week old female Swiss Webster mice were purchased from either Taconic or Charles River, and housed in accordance with the institutional and NIH guidelines on the care and use of animals in research. For ocular HSV infection, mice were anesthetized by intraperitoneal injection of ketamine (100 mg/kg) and xylazine (12 mg/kg). Mice were infected in both eyes by dispensing 10^5^ PFU of HSV1 syn17+ contained in 4 ul following corneal scarification using a 28-gauge needle. For vaginal HSV infection, mice were treated with 2mg of Depo-Provera injected subcutaneously. Five to seven days later, they were anesthetized by intraperitoneal injection of ketamine (100 mg/kg) and xylazine (12 mg/kg) and intravaginally infected with either 5×10^2^ or 10^3^ PFU of HSV1 syn17+ contained in 4 ul using a pipette after clearing the vaginal lumen with a Calginate swab.

For AAV inoculation, mice anesthetized with isoflurane were administered the indicated AAV vector dose by either unilateral intradermal whisker pad (WP), or intravenous injection: unilateral retro-orbital (RO; ocular HSV infected mice) or tail vein (TV; ocular and vaginal HSV infected mice) injection. Tissues were collected at the indicated time.

### Study approval

All animal procedures were approved by the Institutional Animal Care and Use Committee of the Fred Hutchinson Cancer Center. This study was carried out in strict accordance with the recommendations in the Guide for the Care and Use of Laboratory Animals of the National Institutes of Health (“The Guide”).

### HSV reactivation

JQ1 reactivation was performed by intraperitoneal injection of (+)-JQ-1 (JQ1, MedChemExpress) at a dose of 50 mg/kg at the indicated time. JQ1 was prepared from a stock solution (50mg/ml in DMSO, Sigma) by 1:10 dilution in a vehicle solution 10% w/v 2-hydroxypropyl-β-cyclodextrin in PBS (Sigma). Alternatively, JQ1 reactivation was performed with 2 IP injections separated by 12h, which resulted in the detection of virus shedding in the eyes of 6 out of 9 (66.6%) mice (Supplemental figure 5). Hyperthermic shock (HS) reactivation was performed as previously described (Sawtell 2005).

### Cells and herpes simplex viruses

HEK293 (Graham et al. 1977) and Vero cell lines (ATCC # CCL-81) were propagated in Dubelcco’s modified Eagle medium supplemented with 10% fetal bovine serum. HSV-1 strain *syn*17+ (kindly provided by Dr. Sawtell) was propagated and titered on Vero cells.

### AAV production and titering

AAV vector plasmids pscAAV-CBh-m5, and pscAAV-CBh-m8 were used to generate the AAV stocks in this study. All AAV stocks were generated from transfected HEK293 cells and culture media produced by the Viral Vector Core of the Wellstone Muscular Dystrophy Specialized Research Center (Seattle). AAV stocks were generated by PEG-precipitation of virus from cell lysates and culture media, followed by iodixanol gradient separation (Choi et al. 2007; Zolotukhin et al. 1999) and concentration into PBS using an Amicon Ultra-15 column (EMD Millipore). AAV stocks were aliquoted and stored at −80°C. All AAV vector stocks were quantified by qPCR using primers/probe against the AAV ITR, with linearized plasmid DNA as a standard, according to the method of Aurnhammer et al (Aurnhammer et al. 2012). AAV stocks were treated with DNase I and Proteinase K prior to quantification.

### Quantification of viral loads in tissues

Total genomic DNA was isolated from ganglionic tissues using the DNeasy Blood and tissues kit (Qiagen, Germantown, MD) per the manufacturer’s protocol. Viral genomes were quantified by ddPCR in tissue DNA samples using an AAV ITR primer/probe set for AAV, and a gB primer/probe set for HSV as described previously (Aubert et al. 2016). Cell numbers in tissues were quantified by ddPCR using mouse-specific RPP30 primer/probe set: Forward 5′-GGCGTTCGCAGATTTGGA, Reverse 5’-TCCCAGGTGAGCAGCAGTCT, probe 5’-ACCTGAAGGCTCTGCGCGGACTC. In some control ganglia, sporadic samples showed positivity for AAV genomes, although the levels were typically >2-3 logs lower than in ganglia from treated mice having received AAV. We attribute this to low-level contamination of occasional tissue samples. The ganglionic AAV loads for experiments presented in Figure 1 are shown in Supplemental Figure 1 and those for experiments presented in Figures 2, 4, 5, and 6 are shown in Supplemental fig. 6. The statistical analysis was performed using GraphPad Prism version 9.4.1. The test used for each data set is indicated in the figure legends.

**Figure 1:**
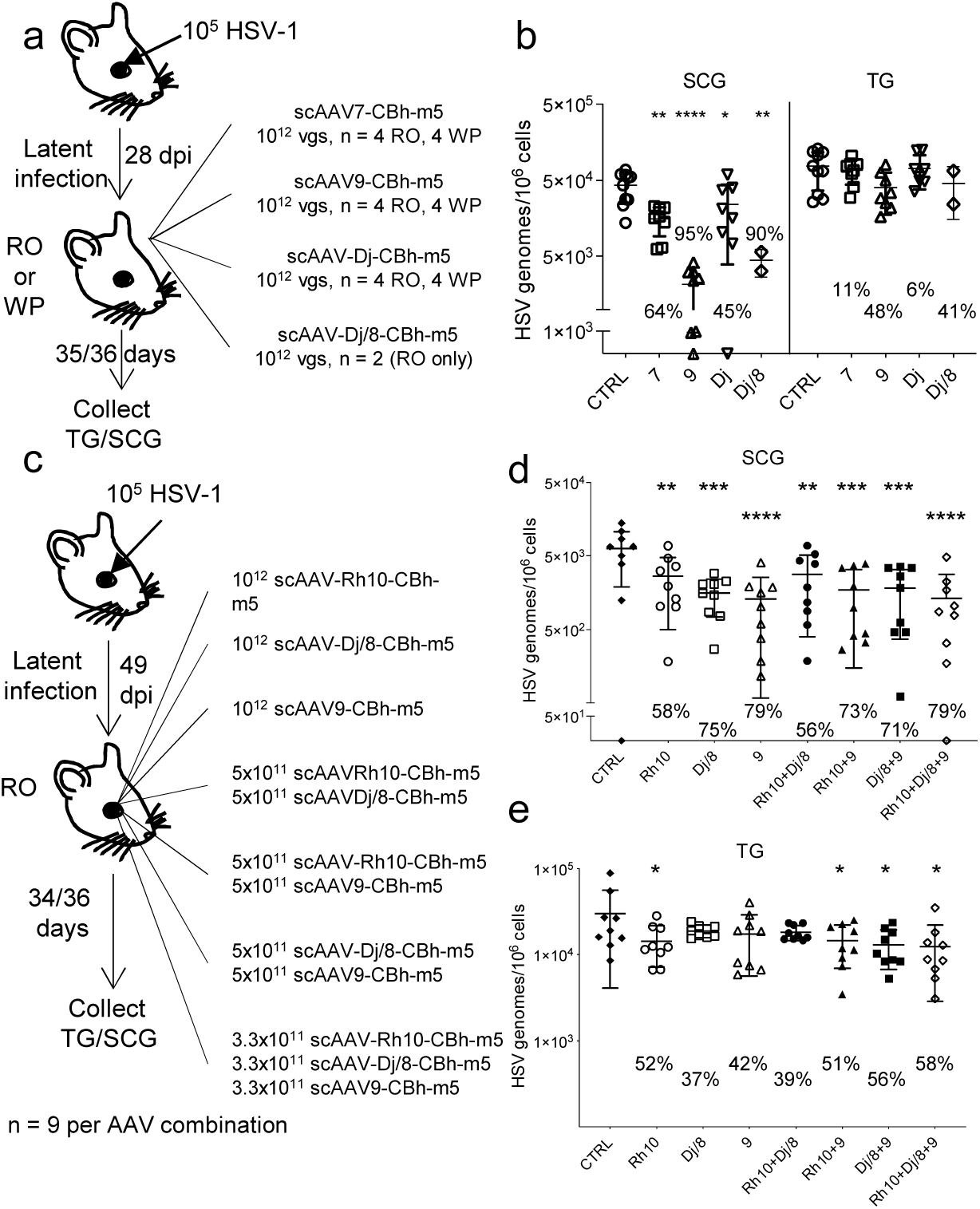
Reduction of ganglionic latent HSV loads after meganuclease therapy delivered using various AAV serotypes. **a**. Experimental timeline of ocular infection and meganuclease therapy. **b**. HSV loads in SCGs and TGs from infected control (CTRL, circles) and infected mice treated with m5 delivered using AAV serotype 7 (squares), 9 (upward triangles), Dj (downward triangles), Dj/8 (diamonds) administered by retro-orbital (RO) or whisker pad (WP) injections at a dose of 10^12^ vg AAV. The percent decrease of HSV loads in treated mice compared to control mice and significant statistical analysis with ordinary one-way Anova, multiple comparisons with *: *p* < 0.05; **: *p* < 0.01, ****: *p* < 0.0001 are indicated for the relevant AAV serotypes. **c**, Experimental timeline of ocular infection and meganuclease therapy. **d**-**e**. HSV loads in SCGs (**d**) and TGs (**e**) from infected control (CTRL, black diamonds) and infected mice treated with m5 delivered using either single AAV serotype Rh10 (open circles) Dj/8 (open squares), 9 (open triangles), dual AAV serotype combinations Rh10+Dj/8 (black circles), Rh10+9 (black triangles), Dj/8+9 (black squares) or triple AAV serotype combination Rh10+Dj/8+9 (open diamonds) administered by RO injection at a total dose of 10^12^ vg AAV. The percent decrease of HSV loads in treated mice compared to control mice and significant statistical difference between treated and control mice obtained with ordinary one-way Anova test with multiple comparisons *: *p* < 0.05; **: *p* < 0.01; ****: *p* < 0.0001 are indicated for the relevant AAV serotype combination. Each graph shows individual and mean values with standard deviation. AAV loads and percent of mutations detected by T7 assay are shown in Supplemental Figure 1.

**Figure 2.**
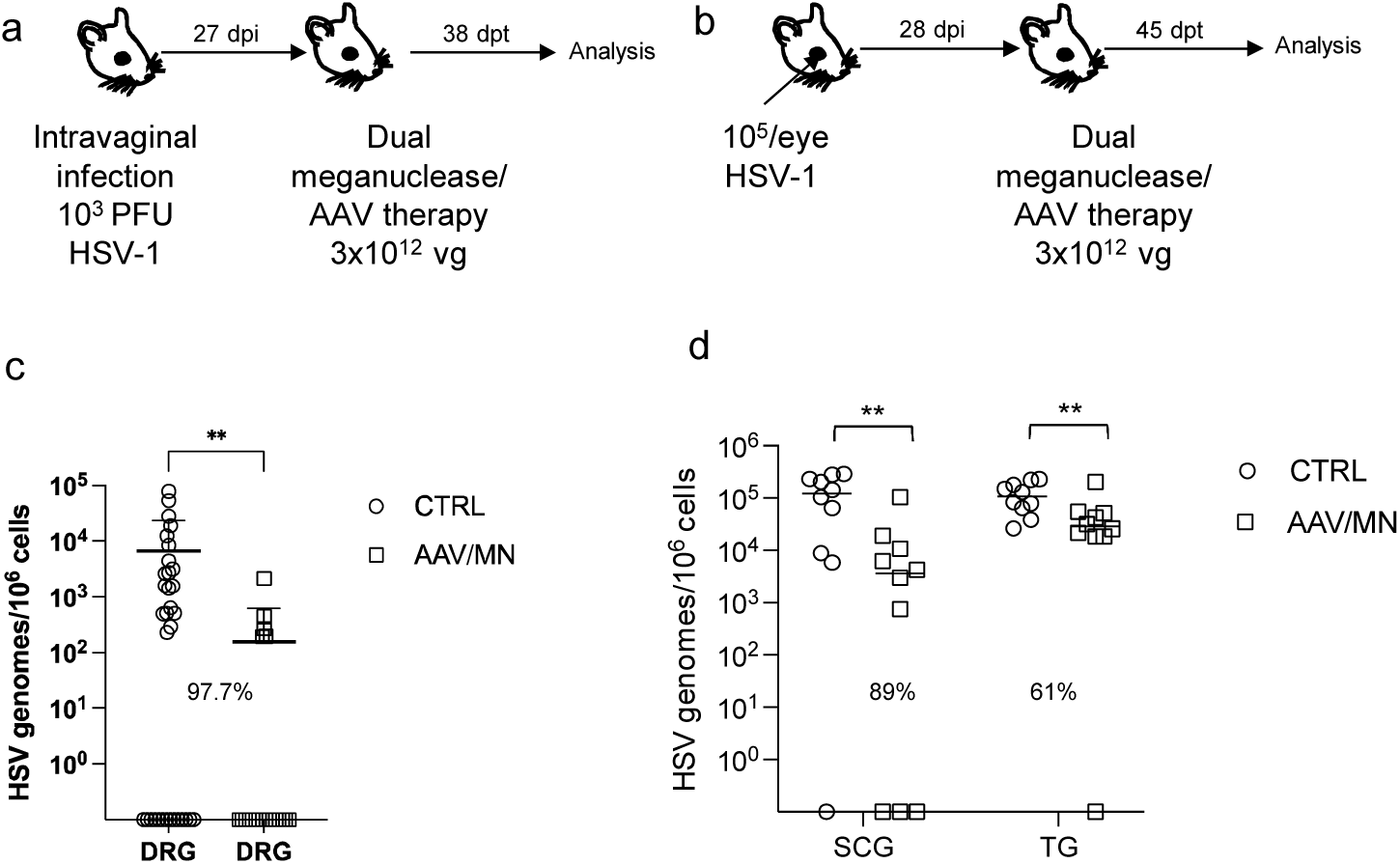
Decrease of ganglionic HSV loads from genital and ocular infection after meganuclease therapy. **a**, Experimental timeline of vaginal infection and meganuclease therapy. **b**, Experimental timeline of ocular infection and meganuclease therapy. **c**. HSV loads in DRGs from control (CTRL n= 7, open circles) and dual meganuclease treated (AAV/MN n = 4, open squares) mice vaginally infected with HSV-1. **d**. HSV loads in SCGs and TGs from control (CTRL n = 10, open circles) and dual meganuclease treated (AAV n = 10, open squares) mice ocularly infected with HSV-1. The percent decrease of HSV loads in treated mice compared to control mice and significant statistical analysis with unpaired one-tailed Mann-Whitney test with **: *p* < 0.01 are indicated. AAV loads are shown in Supplemental figure 7a-b. Each graph shows individual and mean values with standard deviation.

**Figure 3.**
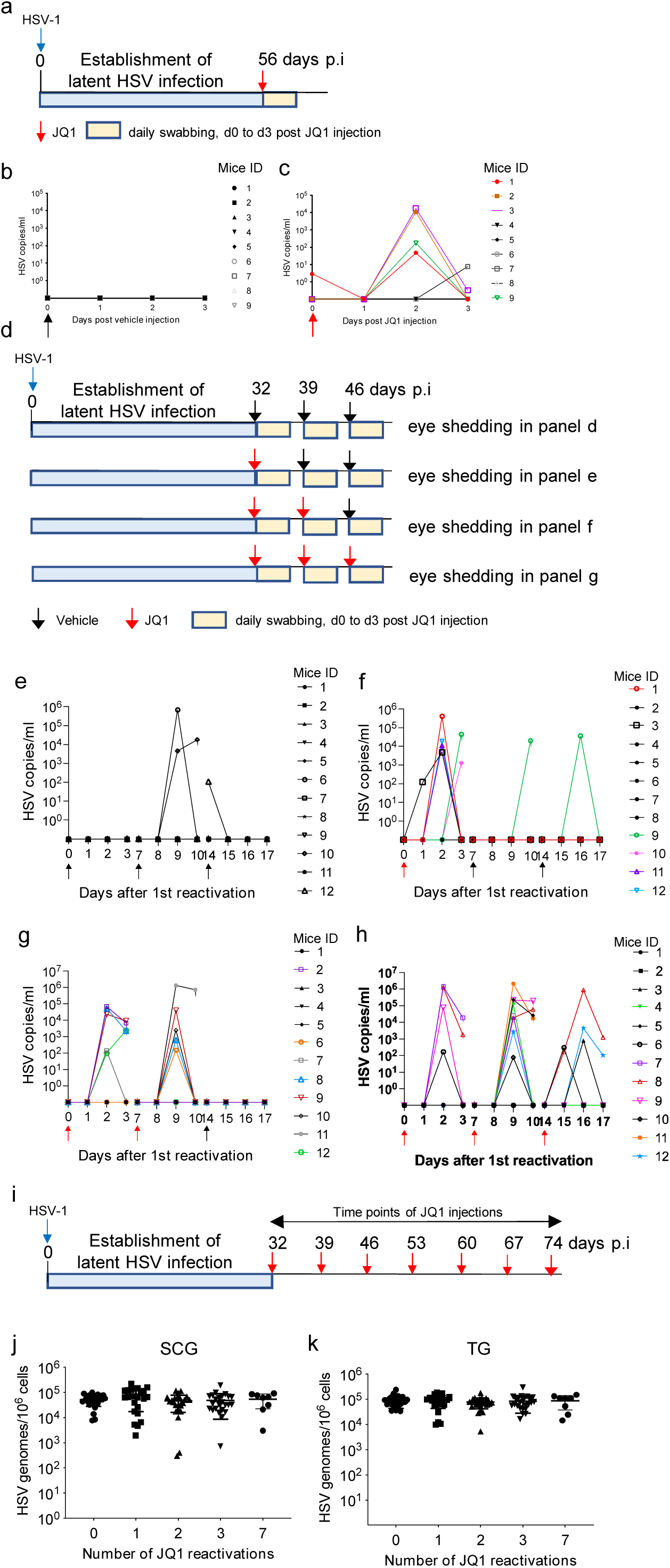
JQ1 reactivation leads to peripheral virus shedding. **a**, Experimental timeline of ocular infection and HSV reactivation. **b**-**c**, Latently infected mice were administered one IP injection of either (**b**) vehicle (n = 9, black arrow), or (**c**) JQ1 (n = 9, red arrow, 50 mg/kg). HSV titers in eye swabs collected from day 0 to 3 post-JQ1 are plotted for each mouse. **d**, Experimental timeline of ocular infection and sequential HSV reactivation with JQ1 injections. **e-h**, HSV titers in eye swabs collected daily after the 1^st^ (day 0), 2^nd^ (day 7) or 3^rd^ (day 14) IP injection of either vehicle (black arrows) or JQ1 (red arrows, 50 mg/kg): **e**, mice (n = 12) received 3 sequential IP injections of vehicle, **f**, mice (n = 12) received 1 IP injection of JQ1 followed by 2 sequential IP injections of vehicle, **g**, mice (n = 12) received 2 sequential IP injections of JQ1 followed by 1 IP injection of vehicle, and **h**, mice received 3 sequential IP injections of JQ1. **i**. Experimental timeline of ocular infection and sequential HSV reactivation with JQ1 injections. **j**-**k**, ddPCR quantification of HSV viral in the SCG (**j**) and TG (**k**) collected from mice having received either 3 sequential injections of vehicle (0, n = 12), 1 JQ1 injection followed by 2 sequential injections of vehicle (1, n = 12), 2 sequential JQ1 injections followed by 1 injection of vehicle (2, n = 12), 3 sequential JQ1 injections (3, n = 12) or 7 sequential JQ1 injections (7, n = 4). Each graph shows individual and mean values with standard deviation.

**Figure 4.**
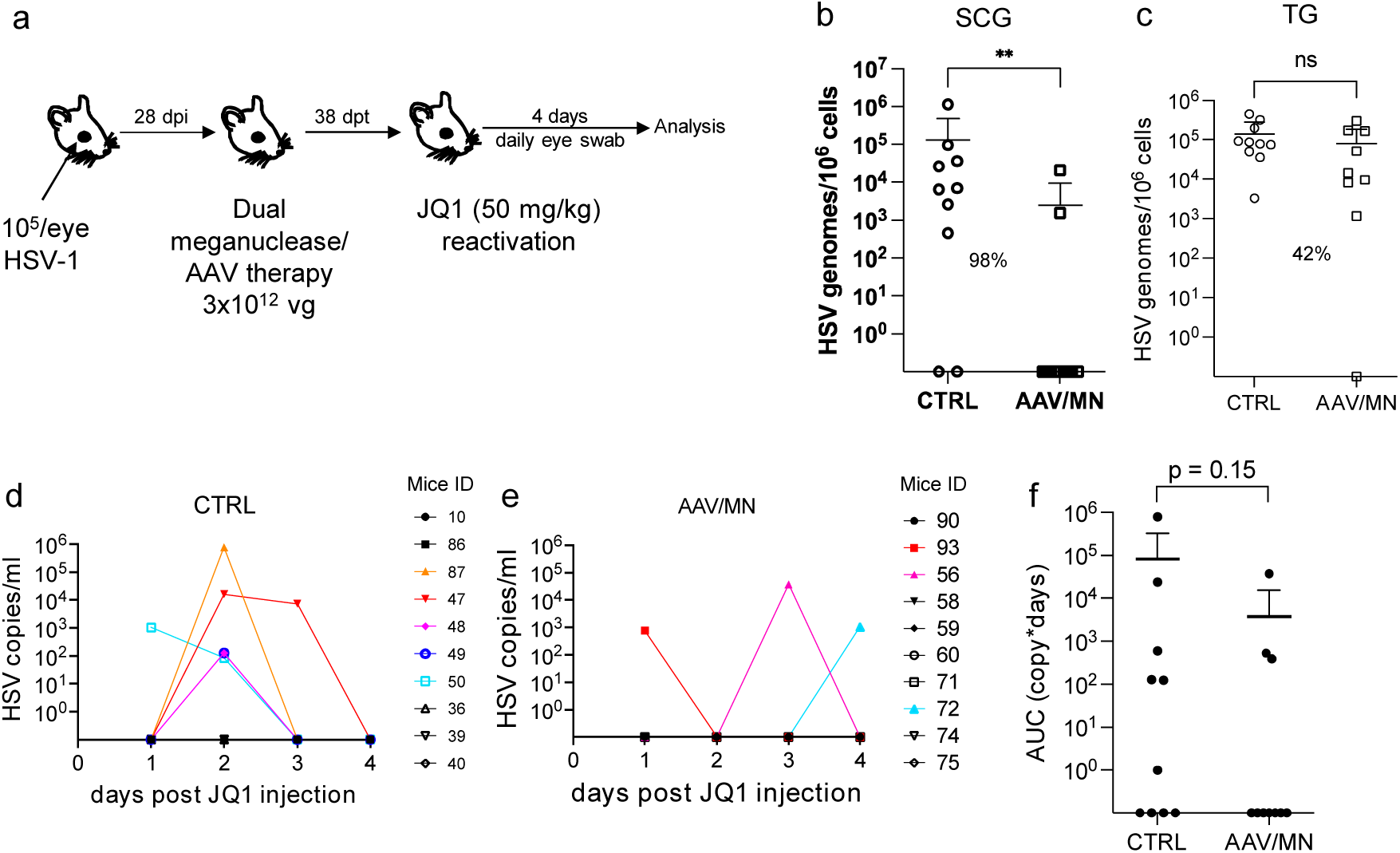
Decrease of peripheral virus shedding in meganuclease-treated mice. **a**, Experimental timeline of ocular infection, meganuclease treatment and viral reactivations with JQ1. **b-c**, HSV loads in SCGs (**b**) and TGs (**c**). The percent decrease of HSV loads in treated mice (n = 10) compared to control mice (n = 10) and statistical analysis using unpaired one-tailed Mann-Whitney test with ns: not significant, **: *p* < 0.01 are indicated. **d-e**. HSV titers in eye swabs collected daily from day 1 to 4 post JQ1 reactivation from control (**d**) and dual meganuclease-treated (**e**) infected mice. **f**, Area under the curve (AUC) analysis with *p* value obtained from statistical analysis using unpaired one-tailed Mann-Whitney test. AAV loads are shown in Supplemental Figure 7c-d. Each graph shows individual and mean values with standard deviation.

**Figure 5.**
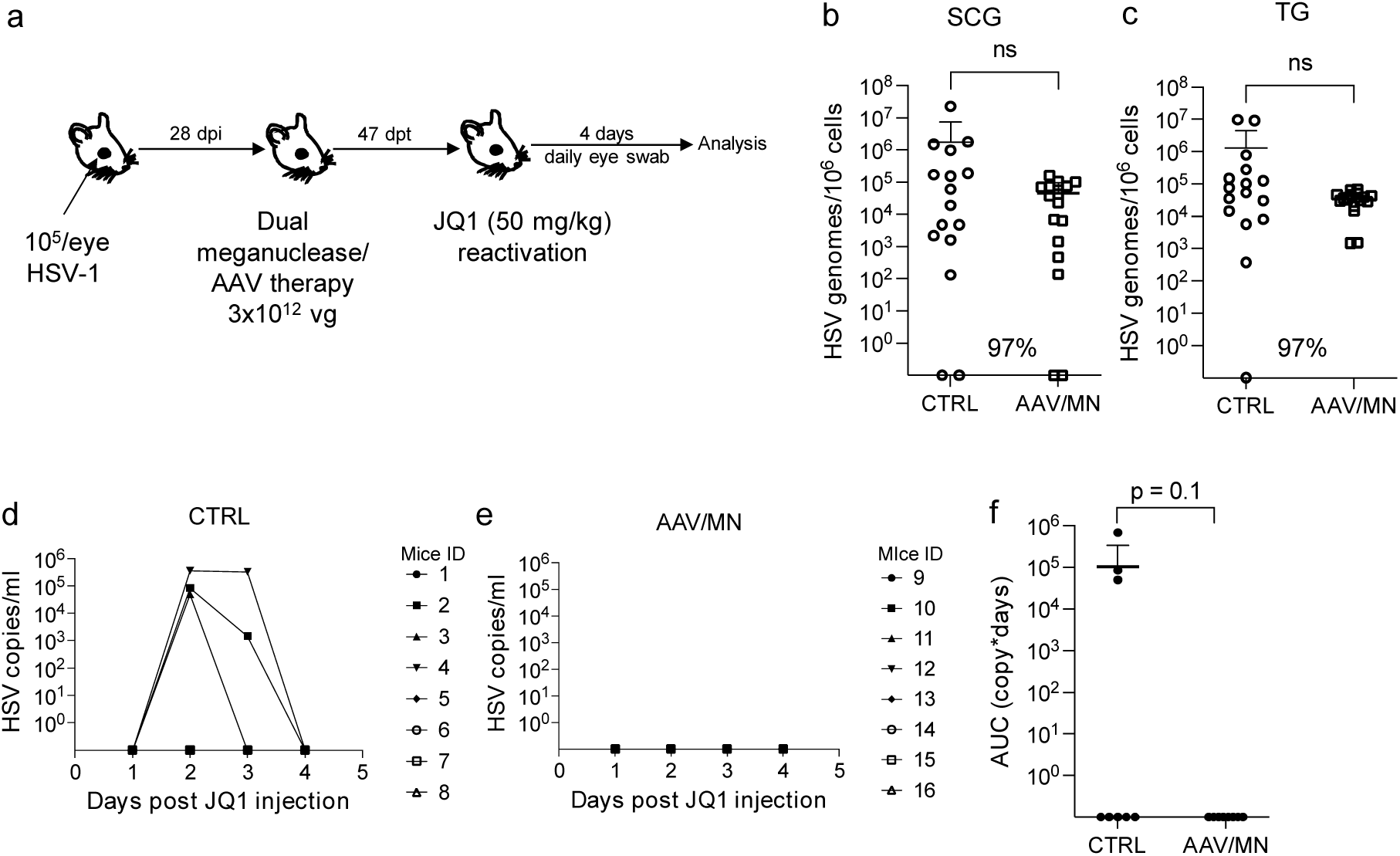
Decrease of peripheral virus shedding in meganuclease-treated mice. **a**, Experimental timeline of ocular infection, meganuclease treatment and viral reactivations with JQ1. **b-c**, HSV loads in SCGs (**b**) and TGs (**c**). The percent decrease of HSV loads in treated mice (n = 8) compared to control mice (n = 8) and statistical analysis using unpaired one-tailed Mann-Whitney test with *: ns not significant are indicated. **d-e**, HSV titers in eye swabs collected daily from day 1 to 4 post JQ1 reactivation from control (**d**) and dual meganuclease-treated (**e**) infected mice. **f**, Area under the curve (AUC) analysis with *p* value obtained from statistical analysis using unpaired one-tailed Mann-Whitney test. AAV loads are shown in Supplemental Figure 7e-f. Each graph shows individual and mean values with standard deviation.

**Figure 6.**
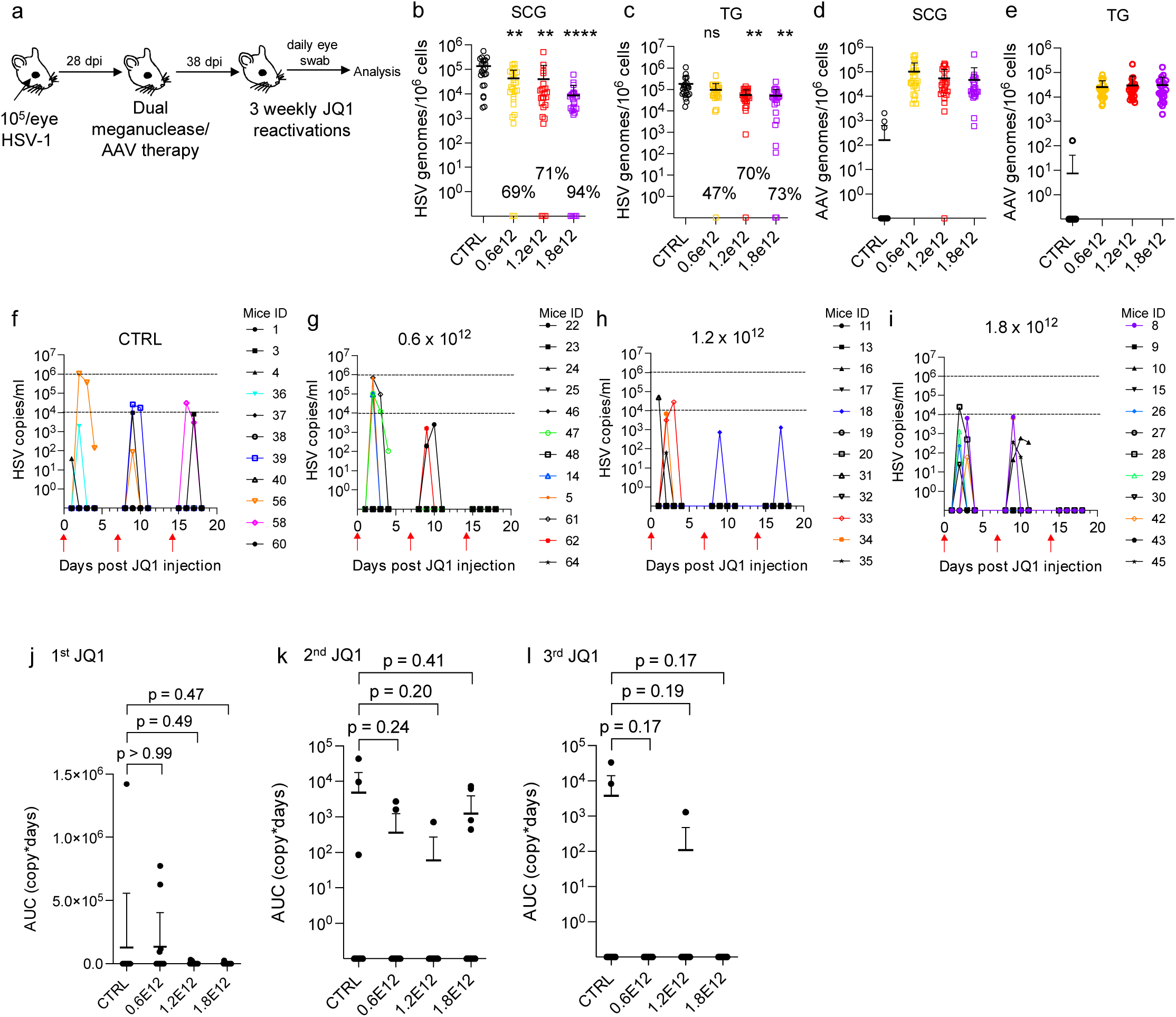
Decrease of peripheral virus shedding in dual meganuclease-treated mice. **a**, Experimental timeline of ocular infection, meganuclease treatment and viral reactivations with JQ1. **b**-**c**, HSV loads and **d**-**e**, AAV loads in SCGs (**b, d**) and TGs (**c, e**) in control infected mice (CTRL n = 11, black circles) and infected mice treated with dual therapy delivered with 0.6 (n = 12, yellow circles), 1.2 (n = 12, red circles) or 1.8 (n = 12, purple circles) ×10^12^ total vg AAV dose. The percent decrease of HSV loads in treated mice compared to control mice and statistical analysis using ordinary one-way Anova, multiple comparisons with ns: not significant, **: *p* < 0.01, ****: *p* < 0.0001 are indicated. **f**-**i**, Virus titers in eye swabs collected daily from day 1 to 4 after each weekly JQ1 reactivation from control infected mice (CTRL, **f**) and infected mice treated with dual therapy delivered with 0.6 (**g**), 1.2 (**h**) or 1.8 (**i**) ×10^12^ total vg AAV dose. **j**-**l**, Area under the curve (AUC) analysis of virus shedding after the first (**j**), the second (**k**), and third (**l**) JQ1 reactivation. The *p* values obtained with the statistical analysis using unpaired, ordinary one-way Anova, with multiple comparisons comparing virus shedding between treatment groups and the control group are indicated. Each graph shows individual and mean values with standard deviation.

### Quantification of HSV in eye swabs

Swab samples were collected into vials containing 1 ml of digestion buffer (KCL, Tris HCl pH8.0, EDTA, Igepal CA-630). DNA was extracted from 200 μl of digestion buffer using QiaAmp 96 DNA Blood Kits (Qiagen, Germantown, MD) and eluted into 100 μl AE buffer (Qiagen). Then, 10 μl of DNA was used to setup 30 μl real-time Taqman quantitative PCR reactions. The primers and probes were as described previously (Jerome et al. 2002). QuantiTect multiplex PCR mix (Qiagen) was used for PCR assays. The PCR cycling conditions were as follows: 1 cycle at 50°C for 2 minutes, 1 cycle at 95°C for 15 minutes, and 45 cycles of 94°C for 1 minute and 60°C for 1 minute. Exo internal control was spiked into each PCR reaction to monitor inhibition. A negative result was accepted only if the internal control was positive with a cycle threshold (CT) within 3 cycles of the Exo CT of no template controls.

### Inflammatory cell foci (ICF) quantification

Liver tissues were paraffin-embedded, sectioned and H&E stained by the Experimental histopathology shared resources of the Fred Hutchinson Cancer Center. ICF were counted by a blinded observer and expressed as the number of ICF per surface area which was determined using Fiji (Schindelin et al. 2012).

### Grading of neuronal changes within trigeminal ganglia

Trigeminal ganglia were paraffin-embedded, sectioned, and H&E stained by the Experimental histopathology shared resources of the Fred Hutchinson Cancer Center. Microscopic changes were graded as to severity by a veterinarian pathologist using a standard grading system whereby 0 = no significant change, 1 = minimal, 2 = mild, 3 = moderate, and 4 = severe.

### Statistical analysis

Statistical analyses for each individual experiment were performed using GraphPad Prism version 9.4.1 and R. Tests were two-sided and p-values smaller than 0.05 considered significant. The specific test used for each analysis is indicated in the corresponding figure legend.

Meta-analyses were performed on combined data from all experiments (figures 4-8). To assess whether AAV/MN-treated mice shed virus less often than control animals and whether the frequency of viral shedding decreased with the duration of meganuclease therapy, we used generalized linear mixed models (GLMM) describing the probability of viral shedding (with viral shedding defined as an AUC > 0) as a function of therapy duration and dose treated as continuous variables, while adjusting for experiment.

Animal-specific random intercepts were included in the model to capture intra-mice dependencies between observations. We also considered including an interaction term between dose and duration to evaluate whether change in the probability of viral shedding over time was affected by dose (in particular, whether it decreased faster with dose). Association between viral shedding and covariates (dose, therapy duration) are reported as odds ratios (OR). The significance of the association between the probability of viral shedding and dose, therapy duration, and their interactions was evaluated using two-sided Wald tests.

To study the relationship between the quantity of virus shedding and therapy dose and duration, we used linear mixed models (LMM) describing the log10-transformed AUC as a function of therapy dose and duration, treated as continuous variables, while adjusting for experiment. The model included an interaction between dose and therapy duration to assess whether change in AUC over time was impacted by dose (e.g., whether the log10-transformed AUC decreased faster with dose). Animal-specific random intercepts were also included to capture intra-mice dependencies between observations. The standard errors of regression coefficients were estimated using a robust, sandwich-type estimator. Association between viral shedding and covariates (dose, therapy duration) are reported as odds ratios (OR). The significance of the association between the log10-transformed AUC and dose, therapy duration, and their interactions was evaluated using Wald tests. Both sets of analyses also evaluated models that included dose squared and square root of dose to perform sensitivity analysis and assess whether the relationship between the probability of viral shedding (or the log10-transformed AUC) and dose were nonlinear. Likewise, models were considered that included squared therapy duration or square root of therapy duration to perform sensitivity analysis and assess whether relationships between the probability of shedding (or the log10-transformed AUC) and therapy duration were nonlinear.

## RESULTS

### Meganuclease therapy reduces ganglionic viral load after ocular or genital HSV infection

We previously evaluated multiple AAV serotypes for delivery of meganucleases to latently-infected mice, and found the best results with AAV-Rh10, followed by AAV8 and AAV1 (Aubert et al. 2020). In an attempt to improve overall gene editing efficiency, we tested additional AAV serotypes that have been reported to be neurotropic including AAV7, AAV9, AAV-DJ, and AAV-DJ/8,(Jacques et al. 2012; Samaranch et al. 2013). Using our previously described model of orofacial HSV disease using ocular infection, we tested each serotype for delivery of a single anti-HSV meganuclease, m5, at an AAV dose of 10^12^ vector genomes (vg) per mouse (Fig. 1a).

Both AAV9 and AAV-Dj/8 administered at a dose of 10^12^ vg were superior to 10^12^ vg AAV-Rh10, the best of our previously-used serotypes (Aubert et al. 2020), showing HSV reductions in superior cervical ganglia (SCG) of 95% (*p* = 10^−5^) and 90% (*p* = 0.018), respectively, relative to untreated controls (Fig. 1b), which compared favorably with the 65% reduction we previously obtained delivering m5 alone with AAV-Rh10 (Aubert et al. 2020). Similarly, both AAV9 and AAV-Dj/8 showed better activity than AAV-Rh10 in trigeminal ganglia (TG), with HSV load reductions of 48% (*p* = 0.07) and 41% (*p* = 0.5), respectively (Fig. 1b), compared with our prior observation of no detectable reduction using AAV-Rh10 to deliver a single meganuclease (Aubert et al. 2020). The route of AAV administration (intravenously via the retro-orbital vein vs. intradermally into the whisker pad) did not have any detectable impact on either AAV transduction or gene editing efficiencies (Supplemental Fig. 1c-e).

Our previous work demonstrated that gene editing of HSV could be increased by using combinations of AAV serotypes for meganuclease delivery, rather than a single AAV serotype, a finding we ascribed to the heterogeneity of neuronal subsets within HSV-infected ganglia (Aubert et al. 2020). We therefore evaluated gene editing with the anti-HSV meganuclease m5 when delivered using single AAV serotypes vs. combinations of AAV9, AAV-Dj/8, and AAV-Rh10 (Fig. 1c) administered as a total dose of 10^12^ vg per mouse. In agreement with our previous results, using combinations of AAV serotypes improved HSV gene editing, with the best efficacy observed using the triple combination of AAV9, AAV-Dj/8, and AAV-Rh10 across both SCG and TG (Fig. 1d-e).

Our previous studies focused on the use of AAV-delivered meganucleases to reduce latent HSV loads in TG and SCG, which are major sites of HSV latency after orofacial infection. However, while orofacial infections with HSV are extremely common, genital infections also represent a major cause of morbidity. To extend our findings to genital HSV, we established latent genital infections in mice by intravaginal inoculation with HSV-1. After establishment of latency, mice were treated with the triple AAV serotype combination (AAV9, AAV-Dj/8, and AAV-Rh10). We chose to deliver therapy using two meganucleases simultaneously (m5+m8), based on our previous work showing that such dual-meganuclease therapy led to improved antiviral efficacy (Aubert et al. 2020). Each animal received a total dose of 3×10^12^ vg AAV (Fig. 2a). In parallel, we tested the same AAV dose against latent orofacial HSV infection as described above (Fig. 2b).

Remarkably, the efficacy in the vaginal model of infection was the highest we have observed to date, with a 97.7% reduction in HSV viral load in the dorsal root ganglia (DRG) where HSV latency is established (Fig. 2c). This compared favorably with the orofacial infection group treated in parallel, in which (in agreement with our previous studies) we observed significant reductions of ganglionic HSV loads of 89% in SCG and 61% in TG (Fig. 2d).

### Induction of HSV shedding using the BET bromodomain inhibitor JQ1

Mice generally show little if any reactivation of latent HSV infection, with minimal to no viral shedding at peripheral sites, limiting their utility in cure studies. The BET bromodomain inhibitor JQ1 has been reported to reactivate latent HSV *in vitro* in primary neuronal cultures, and HSV could be detected in the eyes of about one-third of latently-infected mice treated with JQ1 (Alfonso-Dunn et al. 2017). To evaluate the utility of JQ1 for our cure work, we extended these studies to determine the quantitative kinetics of HSV shedding after JQ1 therapy.

A single intraperitoneal (IP) injection of 50 mg/kg JQ1 given to latently infected mice (Fig. 3a) led to detectable shedding from the eyes of 56% (5/9) of animals, compared with 0/9 animals treated with vehicle alone (Fig. 3b-c). Viral shedding was transient, peaking at 2 days post JQ1, with maximal viral loads ranging from about 10^2^ to almost 10^6^ copies/swab (Fig. 3c). Our results suggest that JQ1 is a more powerful reactivation stimulus for HSV than hyperthermic stress (HS) (Sawtell 2005), which in our hands led to detectable virus shedding in less than 20% of animals (2/12 HS vs 4/10 JQ1), with peak shedding viral loads two logs lower than those observed after JQ1 treatment (Supplemental Fig. 2a-c). Sequential treatment with JQ1 at one-week intervals led to repeated shedding episodes with similar kinetics as observed above (Fig. 3d-h). Over the course of three sequential JQ1 reactivations (Fig. 3h), shedding from individual mice was stochastic; 10/12 (83%) mice shed detectable virus at least once, but only 1/12 (8%) shed after all three treatments, while 4/12 (33%) and 5/12 (42%) shed only after two or one of the three treatments, respectively (Fig. 3h and Supplemental Table 1). Shedding was typically unilateral (only detected in one eye), despite the initial inoculation being to both eyes (unilateral shedding was observed in 33/37 (89%) of events). The side of shedding in one episode was not predictive of the side of future shedding events (Supplemental Table 1). Importantly for cure studies, repeated weekly reactivation of virus with JQ1 up to 7 weekly injections did not change ganglionic viral loads compared with control animals (Fig. 3i-k).

### Reduction of ganglionic HSV load is associated with reduced peripheral shedding

The ability to reproducibly induce HSV reactivation and shedding with JQ1 allowed us to investigate the relationship between ganglionic viral load reduction using meganucleases and subsequent viral shedding at peripheral sites. Latently-infected mice were treated as above using a triple AAV serotype combination (AAV9, AAV-Dj/8, and AAV-Rh10) to deliver the HSV-specific meganucleases m5 and m8 at a total dose of 3×10^12^ vg per animal, or left untreated as controls. One month later, mice from each group were treated with JQ1, and eye swabs were collected daily for 4 days. Ganglionic tissues were then collected to evaluate ganglionic viral loads (Fig. 4a). Consistent with our previous results, ganglionic tissues from treated mice showed a 98% and 42% reduction in mean viral loads in SCG and TG, respectively, when compared to control untreated animals (Fig. 4b-c). After JQ1 administration, only 3/10 (30%) of the dual meganuclease-treated mice had detectable virus in eye swabs, compared with 5/10 (50%) of control untreated animals (Fig 4d-e). The mean titer of HSV in positive eye swabs was 3×10^4^ copies/ml in the meganuclease-treated animals, compared with 1.2×10^5^ copies/ml in the control animals. Area under the curve analysis demonstrated a 95% reduction (*p* = 0.15) in total viral shedding in treated vs. control animals (Fig. 4f). In a separate experiment performed similarly (Fig. 5a), ganglionic tissues from treated mice showed 97% reduction in mean latent HSV genomes in both SCG and TG when compared to control untreated animals (Fig. 5b-c). While 3/8 mice from the control group shed virus with a mean viral titer of 8.2 ×10^5^ copies/ml, 0/8 meganuclease-treated mice had detectable shedding, representing a 100% decrease in total virus shed ((*p* = 0.10), Fig. 5f).

### Safety of AAV/meganuclease therapy

AAV-vectored therapies are generally considered safe. At least 149 human clinical trials are underway or have been completed (Kuzmin et al. 2021), and three AAV-based therapies have received approval for clinical use by the US Food and Drug Administration (FDA) or the European Medicines Agency (EMA). Nevertheless, dose-limiting liver toxicity has been observed after AAV administration in humans, non-human primates, and mice, typically at doses of 2×10^14^ vg/kg or greater. AAV-associated liver toxicity can be ameliorated by steroid administration, and therefore clinically-approved AAV therapeutics such as Zolgensma are typically given after steroid pretreatment (Mendell et al. 2021). The 3×10^12^ vg/animal dose used in the experiments described in Figs. 2, 4 and 5 approached the level associated with liver toxicity in previous studies (mice in our studies received approximately 1×10^14^ vg/kg, assuming an average mouse weight of 30 g), and our animals did not receive steroids. Across multiple studies we observed that 7/70 animals treated with the 3×10^12^ vg/animal dose exhibited clinical signs consistent with hepatotoxicity, including weight change, bloating, and general health decline.

Hepatotoxicity was confirmed in these animals by subsequent histopathological evaluation (Supplemental Fig. 3 and Supplemental Table 2). We therefore evaluated lower total doses of triple AAV serotype/dual meganuclease therapy (0.6, 1.2 or 1.8×10^12^ vg/animal or 1.8, 3.6, or 5.4×10^13^ vg/kg) for their tolerability and effects on viral load and JQ1-induced HSV shedding (Fig. 6a). These doses showed substantially improved tolerability, both clinically and upon histopathological examination and quantification of the number of inflammatory cell foci (ICF) in livers (Supplemental Fig. 4a). Dose-dependent reductions in ganglionic HSV loads were observed across the three treatment groups compared to controls, ranging from 69% and 47% in SCG and TG, respectively, at the 0.6×10^12^ dose to 94% and 73% at the 1.8×10^12^ dose (Fig. 6b-c). To evaluate the effect of these reduced doses on HSV shedding, treated mice were subjected to three weekly rounds of JQ1 administration, followed by eye swabbing as described above. While the percentage of dual meganuclease treated animals shedding virus after the first JQ1 reactivation was not reduced compared with the control mice, it was substantially lower than controls at all doses by the third JQ1 reactivation (0% (0/12), 8% (1/12) and 0% (0/12) for 0.6, 1.2 and 1.8×10^12^ vg/mouse groups, respectively, versus 18% (2/11) in the control group) (Fig. 6f-i). This finding that may relate to the two additional weeks available for meganuclease expression and gene editing activity by the time of the third JQ1 reactivation. Consistent with this interpretation, the reduction in total viral shedding, as determined by AUC analysis, appeared to become more complete over time, with up to a 97-100% reduction in all three groups by the final JQ1 reactivation (Fig. 6j-l). The efficacy of reduced-dose dual meganuclease therapy (1.8×10^12^ vg) was confirmed in a separate experiment (performed identically, except for 7 extra days of AAV-meganuclease therapy prior to only two reactivations with JQ1, Fig. 7a), showing a significant decrease in ganglionic viral loads in both SCG and TG (Fig. 7b-c). In this experiment, 7 of 12 control animals showed detectable viral shedding after JQ1 reactivation, compared with only 1 of 12 animals treated with AAV-meganuclease therapy, (Fig. 7d-e). While none of the treated mice exhibited any clinical signs of hepatotoxicity, we did observe higher numbers of ICF in liver of treated animals receiving the 1.8×10^12^ vg dose compared to control mice (Supplemental Fig. 4b). Histologic analysis of H&E stained TG sections from both control and treated animals revealed subtle evidence of neuronal injury, manifesting as neuronal degeneration, necrosis, and axonopathy. The scores grading prevalence and severity of the microscopic changes were higher in treated animals compared to control mice (Supplemental Fig. 5 and Supplemental Table 3). However, no mice in either the control or experimental group showed detectable signs of neuropathy.

**Figure 7.**
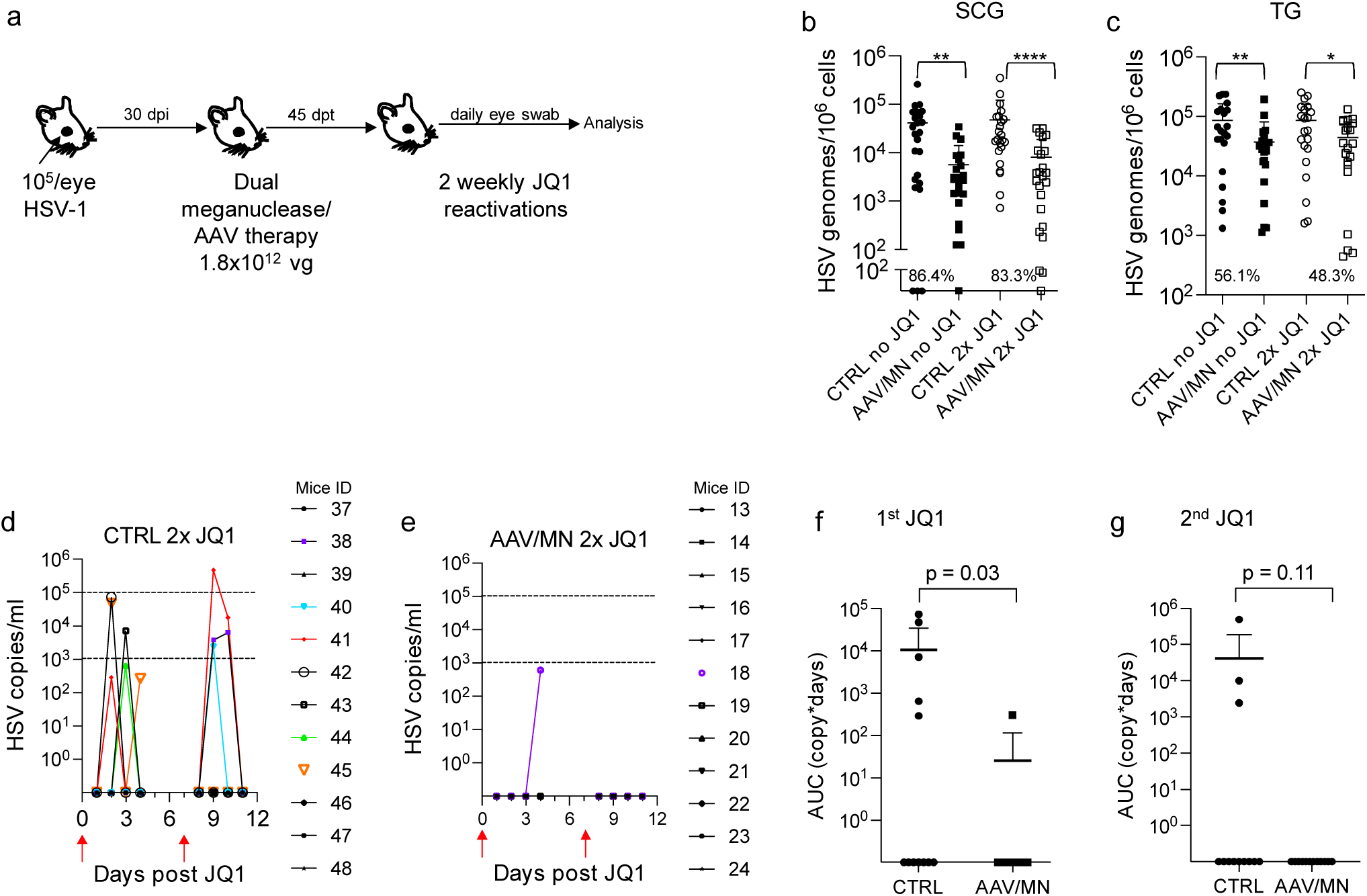
Decrease of peripheral virus shedding in dual meganuclease-treated mice. **a**, Experimental timeline of ocular infection, meganuclease treatment and viral reactivations with JQ1. **b**-**c**, HSV loads in SCGs (**b**) and TGs (**c**) of control infected mice either unreactivated (CTRL no JQ1, n = 12, black circles) or reactivated (CTRL 2x JQ1, n = 12, open circles) and infected mice treated with dual therapy delivered with 1.8 ×10^12^ total AAV dose either unreactivated (AAV/MN no JQ1, n = 12, black squares) or reactivated (AAV/MN 2x JQ1, n = 12, open squares). The percent decrease of HSV loads in treated mice compared to control mice and statistical analysis using unpaired one-tailed Mann-Whitney test with *: *p* < 0.05; **: *p* < 0.01, ****: *p* < 0.0001 are indicated. **d**-**e**, Virus titers in eye swabs collected daily from day 1 to 4 after two weekly JQ1 reactivations (red arrows) from control infected mice (CTRL 2x JQ1, **d**) and infected mice treated with dual therapy delivered with 1.8 ×10^12^ total AAV dose (AAV/MN 2x JQ1, **e**). **f-g**, Area under the curve (AUC) analysis after the first (**f**), and the second (**g**) JQ1 reactivation. The *p* values obtained by statistical analysis with unpaired one-tailed Mann-Whitney test are indicated. AAV loads are shown in Supplemental Figure 7g-h. Each graph shows individual and mean values with standard deviation.

### Reduced dose AAV/meganuclease treatment of vaginally infected animals decreases DRG latent viral load and appears to reduce genital HSV shedding

As noted above, genital HSV infection is a major cause of morbidity in humans. We therefore evaluated the reduced-dose dual meganuclease therapy (total dose of 1.8×10^12^ vg/animal) in vaginally-infected mice (Fig. 8a). The reduced-dose therapy led to a 78.8% (*p* = 0.02) to 95.6% (*p* = 0.006) reduction in latent virus genomes in DRGs (Fig. 8b). Following 3 sequential JQ1 reactivations, 2 out of the 8 control mice shed virus over 2 to 3 sequential days, while only 1 of the 8 AAV-treated mice shed virus, on a single day and at a substantially lower level (Fig. 8c-d). The overall lower rate of reactivation seen in the vaginal model compared to the ocular model may be due to lower levels of ganglionic viral loads in the DRG (10^2^-10^3^ vg/10^6^ cells in DRG, Fig. 8b vs 10^4^-10^5^ vg/10^6^ cells in SCG or TG, Fig. 6b-c).

**Figure 8.**
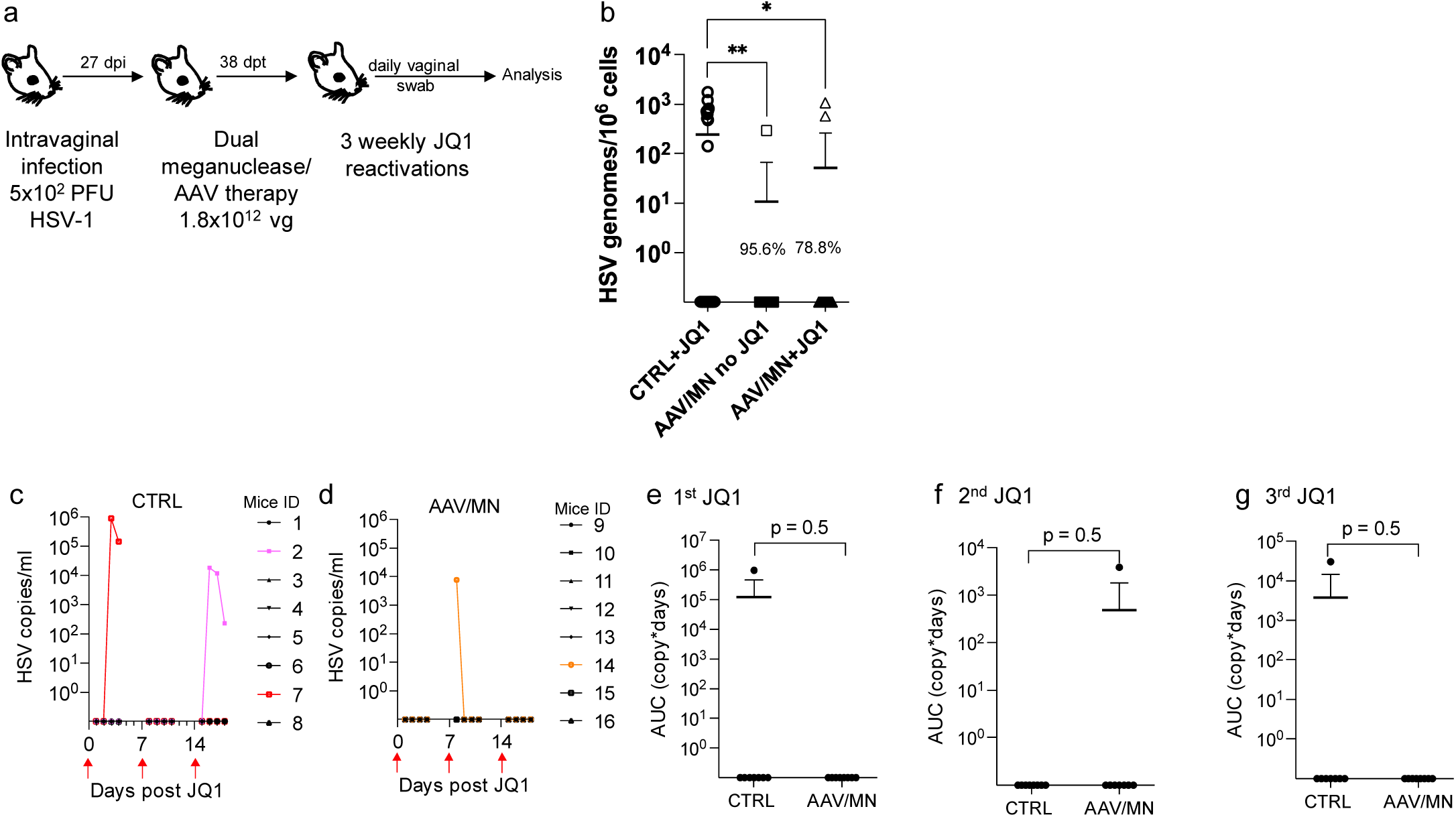
Decrease of genital virus shedding in vaginally infected mice treated with dual meganuclease therapy. **a**, Experimental timeline of intravaginal HSV-1 infection, meganuclease treatment and viral reactivations with JQ1. **b**, HSV loads in DRGs from control infected mice reactivated with 3 weekly JQ1 injections (CTR, n = 8, circles) and infected mice treated with dual therapy unreactivated (AAV/MN no JQ1, n = 8, squares), or reactivated with 3 weekly JQ1 injections (AAV/MN+JQ1, n = 8, triangles). **c-d**. HSV titers in vaginal swabs collected daily from day 1 to 4 post JQ1 injections (indicated by red arrows) from control (CTRL; **c**) and dual meganuclease-treated (AAV/MN; **d**) infected mice. **e**-**h**, Area under the curve (AUC) analysis after the first (**e**), second (**f**), and third (**g**) JQ1 reactivation. The *p* values obtained by statistical analysis with unpaired one-tailed Mann-Whitney test are indicated. The AAV viral loads are shown in Supplemental figure 7i. Each graph shows individual and mean values with standard deviation.

### Meta-analysis of the effect of AAV/meganuclease therapy on HSV shedding

The stochastic nature of HSV reactivation (Schiffer et al. 2009), even when induced by JQ1 (Fig. 3 and Supplemental Table 1), makes evaluation of viral shedding extremely resource-intensive. Practical constraints, including the number of animals that can be studied and housed at the same time, along with the extended duration of each study (∼3 months), limited the statistical power of individual experiments. We therefore performed a meta-analysis of data from all experiments, combining evidence from infection sites (orofacial or genital, Figs. 4-8), thus comparing 174 swabs from AAV/meganuclease-treated animals to 99 swabs from experimentally-matched controls. The primary endpoint was viral shedding, expressed either as a binary variable (equal to 1 for samples in which HSV was detected and 0 otherwise) or the log10-transformed area under the curve (AUC) for quantitative viral shedding. The experiments depicted in Figs. 4-8 represent all of the shedding studies we have performed to date, and each suggest a strong and consistent trend toward a substantial reduction in viral shedding after AAV/meganuclease therapy. The proportion of swabs with detectable HSV was 48% lower among AAV/meganuclease-treated animals compared to controls. The meta-analysis confirmed that animals receiving AAV/meganuclease therapy had a statistically significant reduction in viral shedding (OR = 0.41, *p* = 0.010, GLMM).

We then asked whether dose or duration of meganuclease therapy was associated with the probability of viral shedding (expressed as a binary variable) or the quantity of viral shedding (expressed as the log10-transformed AUC. The data suggest that overall, the probability of viral shedding significantly decreased with the dose of AAV/meganuclease (OR = 0.66; *p* = 0.023, GLMM), and also with the duration of meganuclease therapy (OR = 0.42; *p* < 0.001, GLMM) in treated animals compared to controls. The data further indicate that overall, the quantity of virus shed (AUC) significantly decreased with the AAV/meganuclease dose at a rate of −0.36 log_10_ copy-days per 10^12^ increase in dose (LMM; *p* = 0.028), and also with the duration of meganuclease therapy, at a rate of −0.48 log_10_ copy-days per additional week after treatment (LMM; *p* = 0.017). No significant association was detected between the log10-transformed AUC and the interaction between time and dose (LMM; *p* = 0.59).

## DISCUSSION

Human infection with HSV is lifelong, and while current antiviral approaches can reduce symptoms and transmission, they do not cure. As such, there is a strong and unmet desire among persons infected with HSV for new and potentially curative approaches (Oseso et al. 2016). In the current report, we extend our previous study of gene editing as a potential curative therapy for HSV infection in two important ways. First, we utilize a novel model for small-molecule induction of HSV reactivation in mice to show that a reduction in ganglionic HSV loads of 60-95% via gene editing results in a significant reduction in viral shedding from mice with established orofacial infection. Second, we demonstrate extremely high efficacy of gene editing of latent HSV in DRG after genital HSV infection, with a similar effect on reducing genital HSV shedding. Taken together, these findings address several of the major drivers of interest in HSV cure (Oseso et al. 2016), and strongly support further development of this potentially curative approach for HSV infection.

Mice have proven to be an important laboratory model for many human diseases. Mice are easily infected with HSV, and establish viral latency in a manner that in most ways faithfully reflects human infection. Mouse studies have been critical in defining many aspects of HSV infection, neurovirulence, and immune control (Kollias et al. 2015). A major drawback, however, has been the fact that latent HSV infection in mice exhibits minimal to no spontaneous reactivation and peripheral virus shedding (Hill and Shimeld 1998). Thus, mouse models have been of limited utility in studying HSV therapeutics or vaccines that are directed at control of latent infection and its reactivation. HSV reactivation can be induced *in vivo* by various stimuli such as immunosuppression (Cook et al. 1991), hyperthermic stress (Sawtell and Thompson 1992), or ultraviolet B irradiation (Laycock et al. 1991). These methods show variability in reproducible induction of virus shedding, can be cumbersome, or may be applicable to only certain models that use specific mouse strains or HSV isolates. Here we evaluated the possibility of HSV reactivation by intraperitoneal injection of JQ1, a bromodomain inhibitor that has been proposed as a latency-reversing agent for human immunodeficiency virus (HIV) as part of so-called “kick-and-kill” strategies (Alamer et al. 2021). JQ1 has previously been reported to reactivate HSV in cell culture models of latency (Alfonso-Dunn et al. 2017). Induction of PCR-detectable HSV shedding from the eyes of latently-infected mice has also been reported, although the amount and timing of shedding was not fully defined (Alfonso-Dunn et al. 2017). In this study we demonstrate that JQ1 reproducibly induces detectable viral shedding at the periphery from a substantial fraction of latently-infected mice, at quantitative levels (10^2^ to 10^6^ copies/mL) that are similar to those observed in human observational studies (Ramchandani et al. 2016). Thus, the JQ1 reactivation model should prove extremely useful to the HSV field for future studies regarding the mechanisms and determinants of viral reactivation and peripheral shedding, as wells as interventions such as vaccines or therapeutics aiming to reduce such shedding.

Among people infected with HSV, a major concern, and driver of the desire for cure, is the possibility of transmission of the virus to others (Oseso et al. 2016). While our previous work demonstrated an up to 90% reduction of latent HSV within ganglia after gene editing therapy (Aubert et al. 2020), it remained unclear what effect, if any, such reduction would have on viral shedding at the periphery. The JQ1 reactivation model in mice allowed us to address this issue directly. We found that reduction of ganglionic load via gene editing has a similar effect on viral shedding. Animals receiving AAV/meganuclease therapy showed significant reductions in viral shedding, both in terms of the fraction of samples with detectable virus, and also in the amount of virus shed. While the relationship between HSV shedding quantity and the likelihood of viral transmission remains incompletely understood, previous mathematical modeling suggests that reduction of shedding to levels below 10^4^ viral copies (as observed in most of our treated animals that exhibited residual shedding) would be expected to greatly reduce, if not fully eliminate, the risk of viral transmission (Schiffer et al. 2014).

One challenge in the use of the JQ1 reactivation model in mice is the stochastic nature of the induced viral reactivation and shedding. Within a given cohort of animals, only a subset of mice shed detectable HSV after treatment with JQ1. Upon subsequent JQ1 treatment of the same cohort, a different subset of mice showed detectable shedding, and shedding in one episode was not predictive of subsequent shedding. Similarly, individual mice often shed unilaterally from a single eye; again, this was not predictive of the laterality of subsequent shedding episodes. These findings are similar to observations in humans, in whom shedding is episodic and stochastic, and can occur at distinct anatomical locations during different shedding episodes (Schiffer et al. 2013).

Nevertheless, the fact that only a subset of infected animals shed virus after JQ1 administration constituted a substantial challenge in adequately powering experiments designed to detect not only a reduction in ganglionic viral load, but also a reduction in viral shedding. While each of our individual experiments demonstrated a similar trend toward reduction of HSV shedding after AAV/meganuclease therapy, many individual experiments did not achieve statistical significance. However, meta-analysis of the combined data from all our experiments confirmed a highly significant reduction in HSV shedding from AAV/meganuclease treated animals compared with controls. This reduction in shedding proved to be both dose- and duration of therapy-dependent. The latter observation is particularly encouraging in regard to the potential human translation of the work. While for reasons of practicality we typically evaluate mice for ganglionic load and shedding approximately one month after AAV/meganuclease administration, our data suggests that the efficacy of gene editing continues to increase beyond that time point. Similar continued increases in gene editing efficiency have been reported beyond 100 days in model systems for gene editing of HIV (Wang et al. 2016). In the human setting, a continued improvement in the “completeness” of HSV gene editing at one month or more post meganuclease administration would be expected to provide increasing clinical benefit. Experiments are underway to evaluate the continued improvement in HSV gene editing in mice over periods of a year or more, and to determine whether overall efficacy plateaus at some point.

An ongoing goal of our work is to maximize the efficiency of meganuclease-mediated gene editing, and the subsequent reduction of ganglionic HSV load. We previously demonstrated that both SCG and TG are composed of heterogenous populations of neurons, and that the distribution of both HSV and AAV vector varies between neuronal subsets (Aubert et al. 2020). In our previous work we also demonstrated that a triple AAV combination of AAV1, AAV8, and AAV-Rh10 provided the highest overall gene editing efficiency, and ascribed this to varying tropism of individual serotypes for neuronal subsets (Aubert et al. 2020). Further refinement of our preferred AAV serotypes has led to the AAV9/AAV-Dj/8/AAV-Rh10 triple combination used in this report, which provides marginally better overall gene editing in both SCG and TG compared to our previous combination. Nevertheless, our previous observation holds, in that gene editing within SCG is consistently more efficient than that observed in TG. In this light it is encouraging that our meganuclease therapy led to a significant reduction in orofacial viral shedding, since it remains unclear to what degree the SCG *vs*. TG contributes to peripheral shedding. It is particularly encouraging that gene editing is also highly efficient in reducing latent HSV in DRG, the site of latency for genital HSV infection. As would be predicted from such efficacy, we also observed an apparent reduction in JQ1-induced genital shedding after meganuclease therapy, suggesting that gene editing is also effective against genital disease.

A major issue in human translation of this work is ensuring safety of AAV-delivered meganuclease therapy. AAV vectors in general have been considered quite safe, particularly in comparison with other gene therapy vectors (Bulcha et al. 2021). In addition, to date we have not detected evidence of *in vivo* off-target activity from our meganucleases (Aubert et al. 2016). However, emerging work suggests that AAV vectors may not be as fully innocuous as previously assumed. At high doses AAV vectors can induce liver toxicity, manifesting initially as transaminase elevation. At AAV doses higher than those used in the experiments presented here, liver toxicity can be severe, and has led to liver failure in several animal models (Hinderer et al. 2018). In a particularly tragic example, 3 patients receiving doses of 3×10^14^ vg/kg in a human trial of gene therapy AT132 for X-linked myotubular myopathy (XLMTM) died of fulminant liver failure (Morales et al. 2020). We are therefore encouraged that we observed strong anti-HSV activity at reduced AAV doses that were well tolerated in our mouse model system. Furthermore, in humans liver toxicity can be ameliorated by pretreatment with steroids before AAV therapy (Dasgupta and Keeler 2022). Although we did not evaluate steroid pre-treatment in our mouse studies, the availability of steroid pretreatment and reduced AAV dosing options provides confidence that anti-HSV efficacy can be achieved at well-tolerated AAV doses.

Another newly-appreciated potential AAV toxicity is damage to ganglionic neurons. Histological evidence of neuronal injury after AAV administration has been described in mice, rats, piglets, dogs, and non-human primates (Hinderer et al. 2018; Palazzi et al. 2022; Schuster et al. 2014; Hinderer et al. 2016), as well as in at least two human trial participants at autopsy (Mullard 2021). The causative mechanism of such injury remains unclear; among the current leading hypotheses are saturation of neural protein-folding capacity (Hordeaux, Buza, Jeffrey, et al. 2020) and TLR9-mediated recognition of vector or transgene RNA (Hamilton and Wright 2021). Despite histological evidence of neuronal injury, clinical signs of injury in experimental animals have been rare, consisting mainly of mild gait or balance disturbance (Hinderer et al. 2018; Hordeaux, Buza, Dyer, et al. 2020; Keiser et al. 2021). Such signs have only been reported in a single patient among several thousand human participants in various trials of AAV-delivered gene therapies (Kuzmin et al. 2021). Nevertheless, reports of neuronal toxicity are of particular relevance for our work, as the neuron itself is the cellular site for anti-HSV gene editing. Consistent with other studies, we have observed subtle evidence of neuronal injury in experimental mice, manifesting as neuronal degeneration, necrosis, and axonopathy, although our mice have not shown any associated behavioral alterations. However, mice may not be an ideal model system to evaluate such effects, and future studies in alternative animal models of HSV infection, such as guinea pigs or even non-human primates, may be warranted to ensure maximal safety. If such studies confirm the dramatic anti-HSV effect we have observed in mice, along with an acceptable safety profile, human trials may be warranted.

## ACKNOWLEDGMENTS

The authors are grateful for support provided by R01 AI132599 (Jerome), the Caladan Foundation, over 1600 individual donors, the Viral Vector Core of the Wellstone Muscular Dystrophy Specialized Research Center (Seattle, grant number: 5P50AR065139), and the Fred Hutchinson Cancer Center Shared Resources Division (NIH/NCI Cancer Center Support Grant P30 CA015704). We are also grateful to Cellectis (Paris, France) for the original development of meganucleases m4, m5, and m8, as well as Philippe Duchateau and Roman Galetto (Cellectis) for helpful discussions.

