## Supplemental Figures and Tables for "AAV-delivered gene editing for latent genital or orofacial herpes simplex virus infection reduces ganglionic viral load and minimizes subsequent viral shedding in mice"

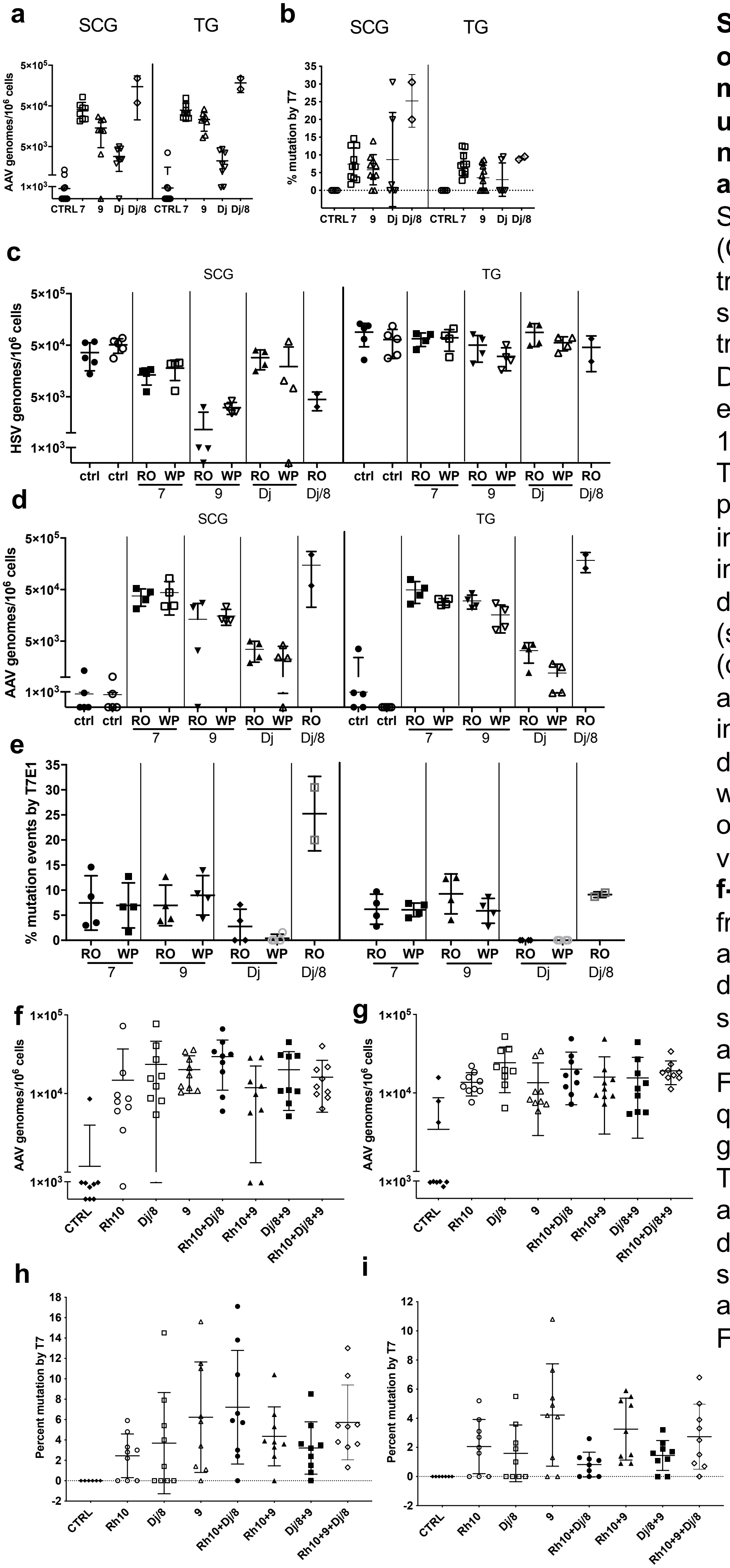

**Supplemental Figure 1. Reduction of ganglionic HSV loads after meganuclease therapy delivered using various AAV serotypes does not depend on the route of administration**. **a**, AAV loads in SCGs and TGs from infected control (CTRL, circles) and infected mice treated with m5 delivered using AAV serotype 7 (squares), 9 (upward triangles), Dj (downward triangles), Dj/8 (diamonds) administered by either RO or WP injections (see Fig. 1a). **b**, Percent mutation quantified by T7 assay in latent HSV genomes present in SCG and TG collected from infected control (CTRL, circles) and infected mice treated with m5 delivered using AAV serotype 7 (squares), 9 (upward triangles), Dj (downward triangles), Dj/8 (diamonds) administered by either RO or WP injections (see Fig. 1a). **c-e**, Same data as above in panels **a** and **b** as well in **Figure 1b** presented per route of administration of the AAV delivery vectors.
**f-g**, AAV loads in SCG (**f**) and TG (**g**) from infected control (CTRL, circles) and infected mice treated with m5 delivered using AAV combination of serotype 9, Dj/8 and Rh10 administered by RO injections (see Fig. 1c). **h-i**, Percent mutation quantified by T7 assay in latent HSV genomes present in SCG (**h**) and TG(**i**) collected from infected control and infected mice treated with m5 delivered using AAV combination of serotype 9, Dj/8 and Rh10 administered by RO injections (see Fig. 1d).

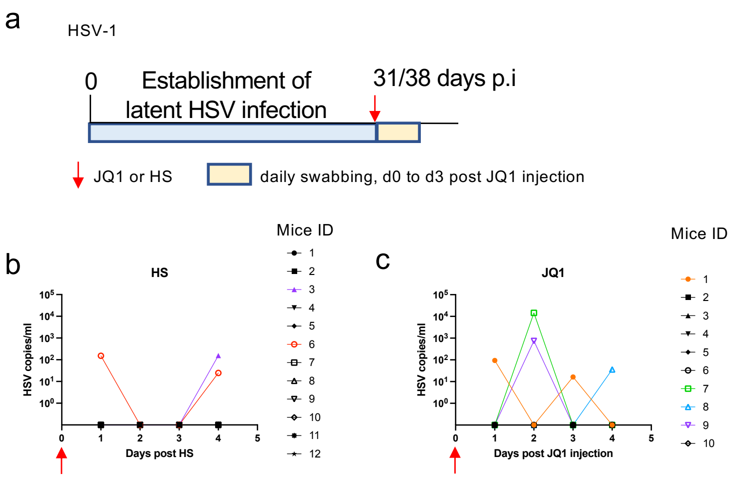

**Supplemental figure 2. Virus shedding after HSV reactivation by hyperthermic stress or JQ1 injection. a**, Experimental timeline of ocular infection and HSV reactivation using hyperthermic stress (31 dpi) or JQ1 injection (38 dpi). **b-c**, Latently infected mice were reactivated at day 0 (red arrow) by either (**b**) hyperthermic stress n = 12, or (**c**) JQ1 IP injection (50 mg/kg) n = 10. HSV titers in eye swabs collected from day 1 to 4 post-JQ1 are plotted for each mouse.

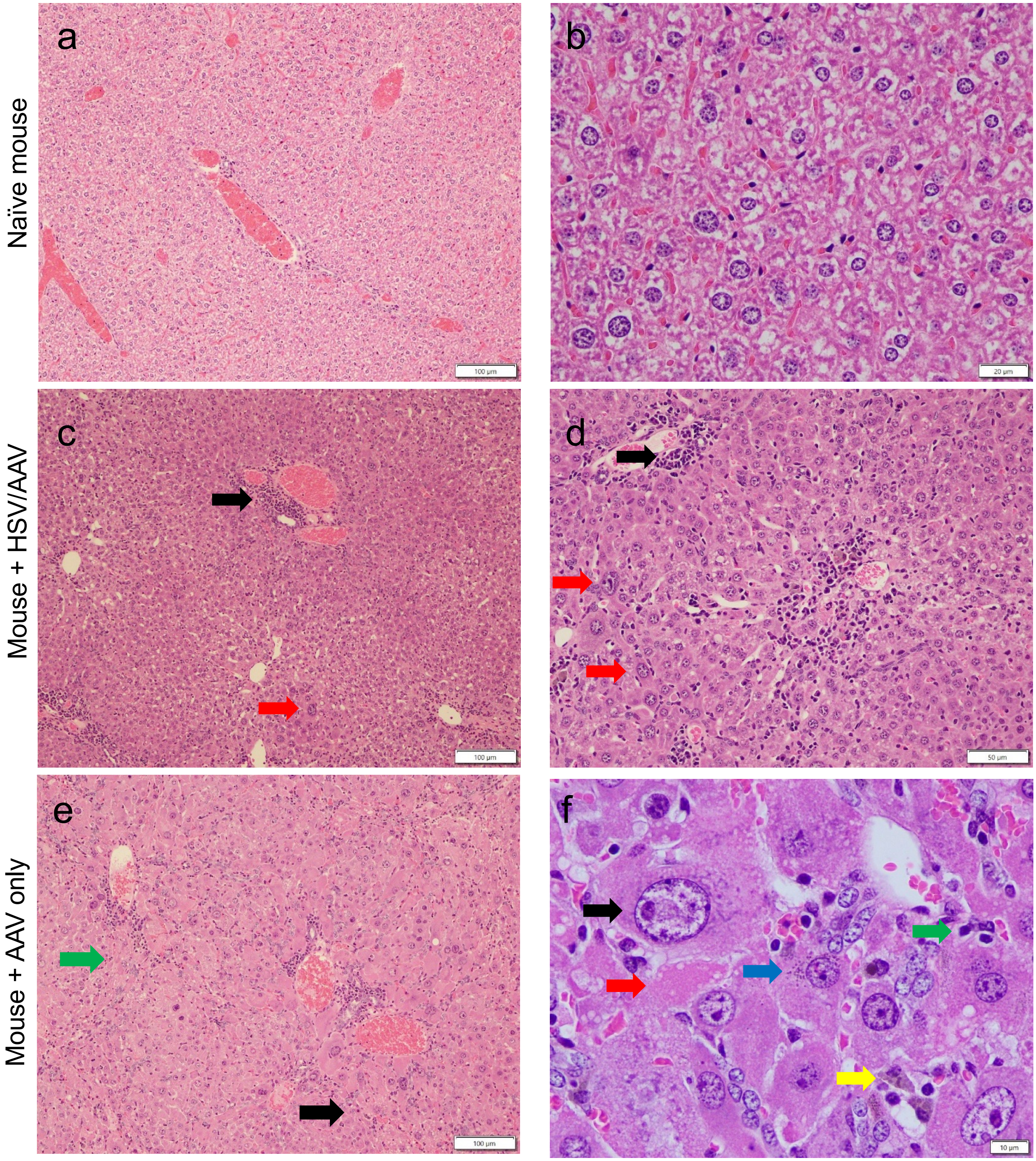

**Supplemental figure 3: Histopathology of liver from dual meganuclease-treated mice**.
H&E staining of liver section from naive mouse (a,10x and b, 40x), HSV-infected mouse administered $3x10^{12}$ vg AAV (c, 10x and d, 40x) and mouse administered $3x10^{12}$ vg AAV only (e, 10x and f, 60x).
Black arrows indicate hepatocellular karyomegaly, anisocytosis and anisokaryosis, red arrows indicate hepatocellular necrosis, green arrows indicate periportal or parenchymal mixed infiltrates, yellow arrow indicates pigmented Kupffer cells, blue arrow indicates suspected biliary cholestasis.

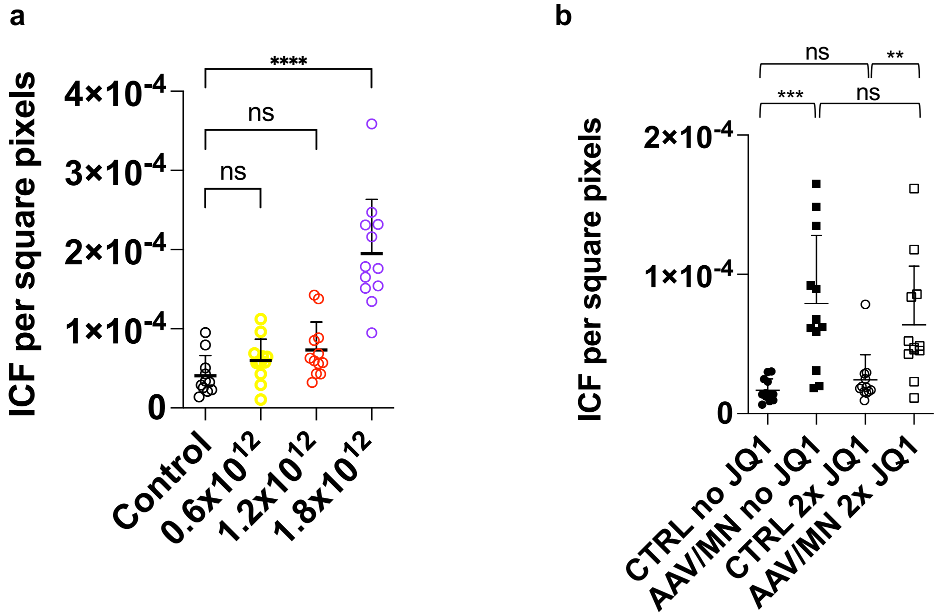

**Supplemental figure 4: Inflammatory cell foci in liver of meganuclease-treated mice. a**, ICF in liver sections from either HSV infected control mice (control, black circles), or treated with dual meganuclease therapy at a dose of $0.6 \times 10^{12}$ vg AAV (yellow circles), $1.2 \times 10^{12}$ vg AAV (red circles) and $1.8 \times 10^{12}$ vg AAV (purple circles) from experiment described in Figure 5a-l. Statistical analysis using ordinary one-way Anova with multiple comparisons test, ns: not significant; ****: $p < 0.0001$. **b**, ICF in liver sections from either HSV infected control mice unreactivated (control no JQ1, black circles), control mice reactivated with JQ1 (control 2x JQ1, black squares), HSV infected mice treated with dual meganuclease therapy unreactivated (AAV/MN no JQ1, open circles), or reactivated with JQ1 (AAV/MN 2x JQ1, open squares) from experiment described in Figure 5m-u. Statistical analysis using unpaired one-tailed $t$ test. ns: not significant; **: $p < 0.01$; ***: $p < 0.001$.

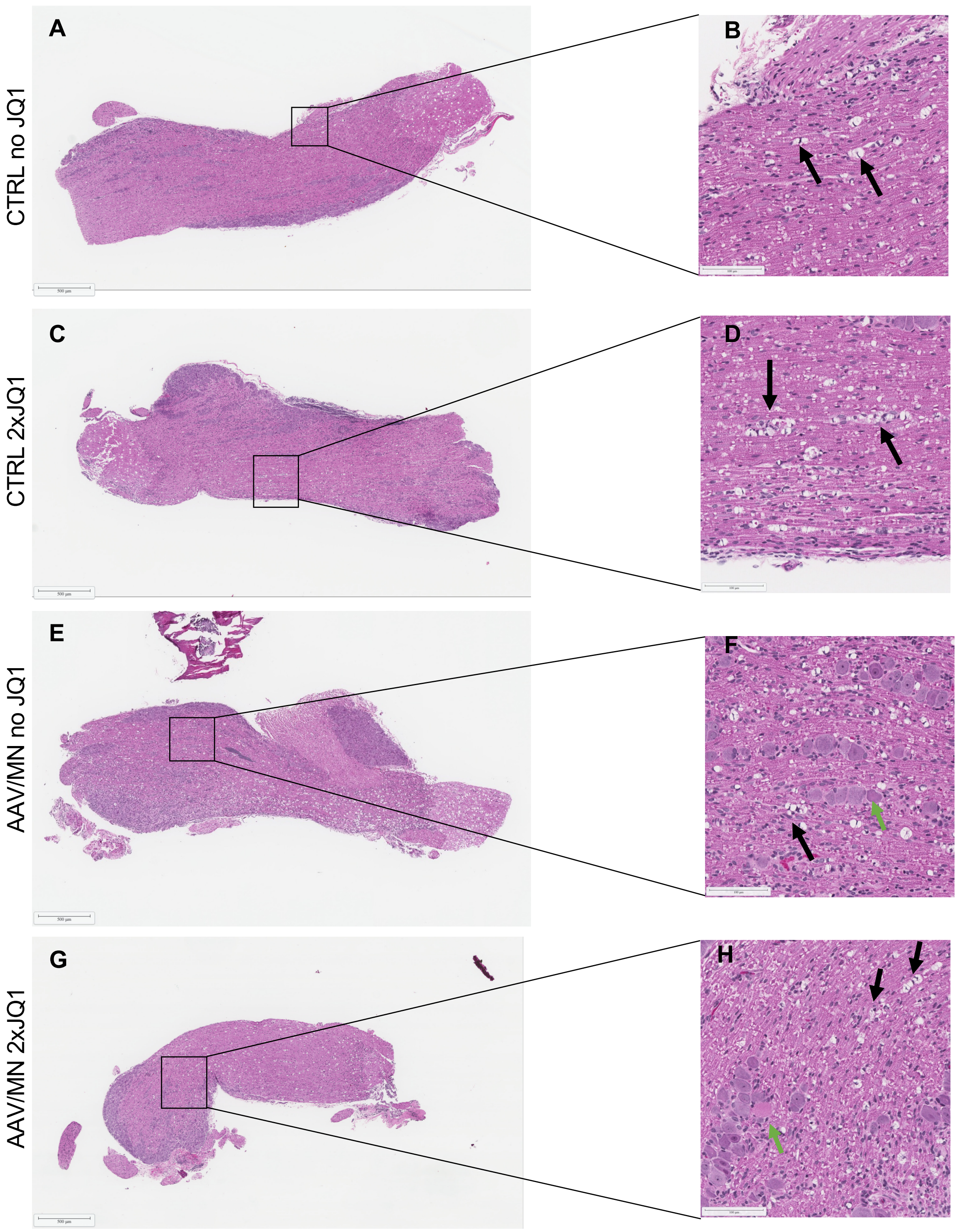

**Supplemental figure 5: Histopathology of TG from dual meganuclease-treated mice**. Images of H&E stained trigeminal ganglia sections from either latently infected control mice not reactivated (CTRL no JQ1 (slide 10 in Supplemental Table 3): **A**, 2.5x and **B**,20x), and reactivated with JQ1 (CTRL 2xJQ1 (slide 11 in Supplemental Table 3): **C**, 2.5x and **D**, 20x) or $1.8x10^{12}$ vg AAV/dual meganuclease treated mice not reactivated (AAV/MN no JQ1 (slide 1 in Supplemental Table 3): **E**, 2.5x and **F**, 20x) and reactivated (AAV/MN 2xJQ1 (slide 4 in Supplemental Table 3): **G**, 2.5x and **H**, 20x). Black arrows indicate signs of axonopathy, green arrows indicate neurons with central chromatolysis.

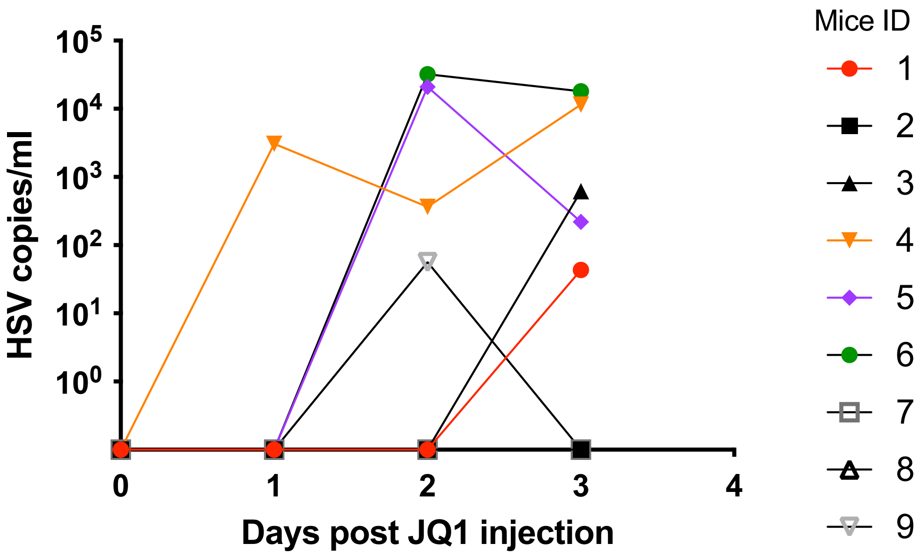

**Supplemental Figure 6. Viral shedding after a double dose of JQ1 in 67% (6 /9) of reactivated mice**. **a**, Latently infected mice were administered 2 IP injections of JQ1 (50 mg/kg) separated by 12h, n = 9. HSV titers in eye swabs collected from day 0 to 3 post-JQ1 are plotted for each mouse.

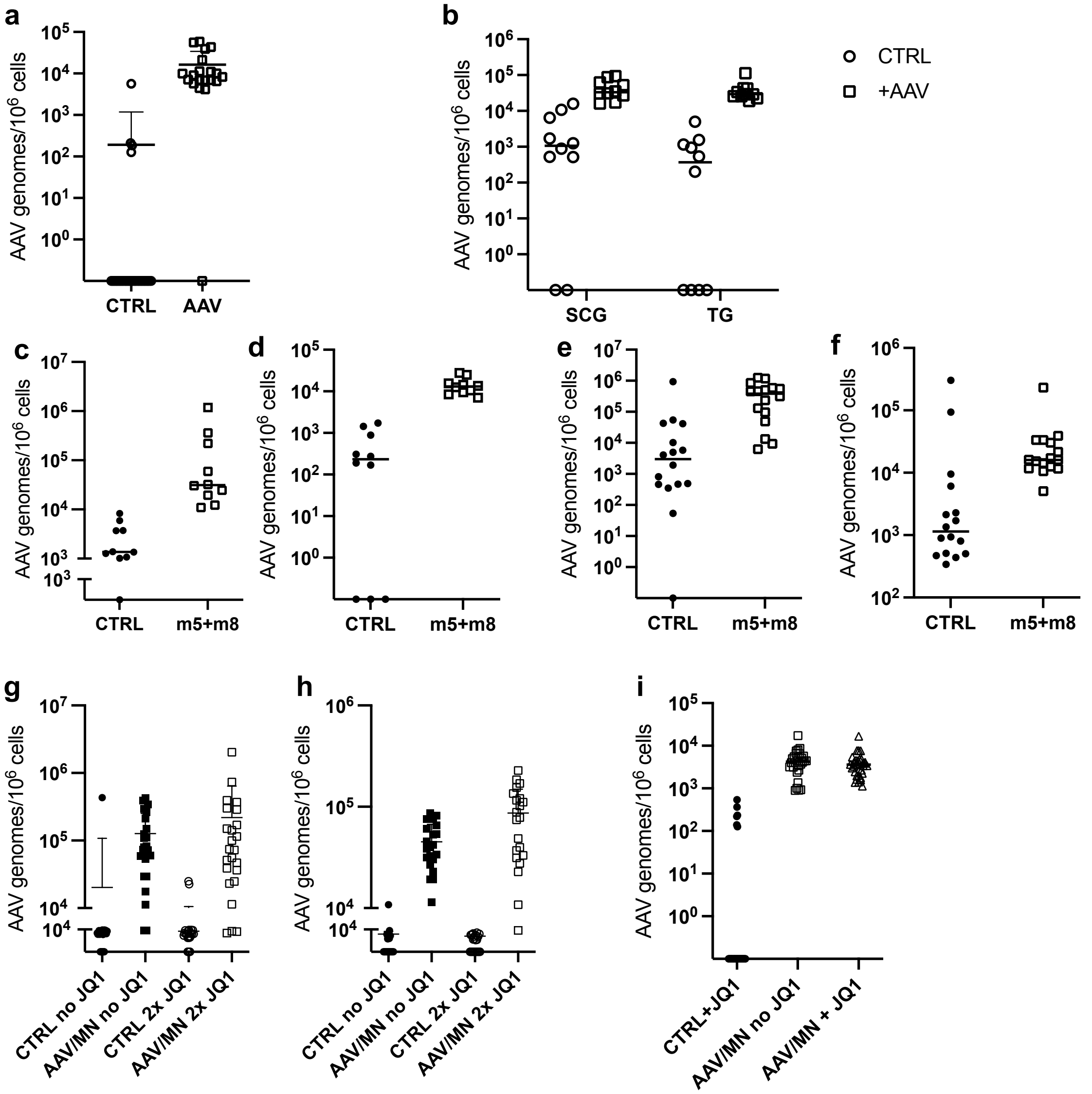

**Supplemental Figure 7. ddPCR quantification of AAV viral loads**. **a**, AAV loads in DRGs from latently infected mice following intravaginal administration of HSV1 either control untreated (CTRL, open circles) or treated with AAV-delivered meganuclease dual therapy (AAV, open squares) in the experiment presented in Figure 2a, c. **b**, AAV loads in SCG and TG from latently infected mice following ocular administration of HSV1 either control untreated (CTRL, open circles) or treated with AAV-delivered meganuclease dual therapy (AAV, open squares) in the experiment presented in Figure 2b, d. **c**-**d**, AAV loads in SCGs (**c**) and TGs (**d**) from latently infected mice following ocular administration of HSV1 either control untreated (CTRL, circles) or treated with AAV-delivered meganuclease dual therapy (AAV, open squares) in the experiment presented in Figure 4a-f. **e**-**f**, AAV loads in SCGs (**e**) and TGs (**f**) from latently infected mice following ocular administration of HSV1 either control untreated (CTRL, circles) or treated with AAV-delivered meganuclease dual therapy (AAV, open squares) in the experiment presented in Figure 4g-l. **g**-**h**, AAV loads in SCGs (**g**) and TGs (**h**) from latently infected mice following ocular administration of HSV1 either control untreatedno reactivated (CTRL no JQ1, black circles), treated with AAV-delivered meganuclease dual therapy not reactivated (AAV/MN no JQ1, black squares), control untreated reactivated twice with JQ1 (CTRL 2x JQ1, circles) or treated with AAV-delivered meganuclease dual therapy reactivated twice with JQ1 (AAV/MN 2x JQ1, open squares) in the experiment presented in Figure 5n-u. **i**, AAV loads in DRGs from latently infected mice following intravaginal administration of HSV1 either control untreated reactivated with JQ1 (CTRL+JQ1, circles), treated with AAV-delivered meganuclease dual therapy not reactivated (AAV/MN no JQ1, open squares), or treated with AAV-delivered meganuclease dual therapy reactivated with JQ1 (AAV/MN+JQ1, open triangles) in the experiment presented in Figure 6.

**Supplemental Table 1.** Virus titers in eye swabs collected after mock and JQ1 reactivations.

#### Group 1[b]: vehicle - vehicle - vehicle

| Mouse | 1 | | 2 | | 3 | | 4 | | 5 | | 6 | | 7 | | 8 | | 9 | | 10 | | 11 | | 12 | |
|---|---|---|---|---|---|---|---|---|---|---|---|---|---|---|---|---|---|---|---|---|---|---|---|---|
| Day/eye[c] | L | R | L | R | L | R | L | R | L | R | L | R | L | R | L | R | L | R | L | R | L | R | L | R |
| 0 | 0 | 0 | 0 | 0 | 0 | 0 | 0 | 0 | 0 | 0 | 0 | 0 | 0 | 0 | 0 | 0 | 0 | 0 | 0 | 0 | 0 | 0 | 0 | 0 |
| 1 | 0 | 0 | 0 | 0 | 0 | 0 | 0 | 0 | 0 | 0 | 0 | 0 | 0 | 0 | 0 | 0 | 0 | 0 | 0 | 0 | 0 | 0 | 0 | 0 |
| 2 | 0 | 0 | 0 | 0 | 0 | 0 | 0 | 0 | 0 | 0 | 0 | 0 | 0 | 0 | 0 | 0 | 0 | 0 | 0 | 0 | 0 | 0 | 0 | 0 |
| 3 | 0 | 0 | 0 | 0 | 0 | 0 | 0 | 0 | 0 | 0 | 0 | 0 | 0 | 0 | 0 | 0 | 0 | 0 | 0 | 0 | 0 | 0 | 0 | 0 |
| 7 | 0 | 0 | 0 | 0 | 0 | 0 | 0 | 0 | 0 | 0 | 0 | 0 | 0 | 0 | 0 | 0 | 0 | 0 | 0 | 0 | 0 | 0 | 0 | 0 |
| 8 | 0 | 0 | 0 | 0 | 0 | 0 | 0 | 0 | 0 | 0 | 0 | 0 | 0 | 0 | 0 | 0 | 0 | 0 | 0 | 0 | 0 | 0 | 0 | 0 |
| 9 | 0 | 0 | 0 | 0 | 0 | 0 | 0 | 0 | 0 | 0 | 0 | 1379357 | 0 | 0 | 0 | 0 | 0 | 0 | 0 | 9240 | 0 | 0 | 0 | 0 |
| 10 | 0 | 0 | 0 | 0 | 0 | 0 | 0 | 0 | 0 | 0 | 0 | 0 | 0 | 0 | 0 | 0 | 0 | 0 | 0 | 36992 | 0 | 0 | 0 | 0 |
| 14 | 0 | 0 | 0 | 0 | 0 | 0 | 0 | 0 | 0 | 0 | 0 | 0 | 0 | 0 | 0 | 0 | 0 | 0 | 0 | 0 | 0 | 0 | 0 | 214 |
| 15 | 0 | 0 | 0 | 0 | 0 | 0 | 0 | 0 | 0 | 0 | 0 | 0 | 0 | 0 | 0 | 0 | 0 | 0 | 0 | 0 | 0 | 0 | 0 | 0 |
| 16 | 0 | 0 | 0 | 0 | 0 | 0 | 0 | 0 | 0 | 0 | 0 | 0 | 0 | 0 | 0 | 0 | 0 | 0 | 0 | 0 | 0 | 0 | 0 | 0 |
| 17 | 0 | 0 | 0 | 0 | 0 | 0 | 0 | 0 | 0 | 0 | 0 | 0 | 0 | 0 | 0 | 0 | 0 | 0 | 0 | 0 | 0 | 0 | 0 | 0 |

#### Group 2[b]: JQ1 - vehicle - vehicle

| Mouse | 13 | | 14 | | 15 | | 16 | | 17 | | 18 | | 19 | | 20 | | 21 | | 22 | | 23 | | 24 | |
|---|---|---|---|---|---|---|---|---|---|---|---|---|---|---|---|---|---|---|---|---|---|---|---|---|
| Day/eye[c] | L | R | L | R | L | R | L | R | L | R | L | R | L | R | L | R | L | R | L | R | L | R | L | R |
| 0 | 0 | 0 | 0 | 0 | 0 | 0 | 0 | 0 | 0 | 0 | 0 | 0 | 0 | 0 | 0 | 0 | 0 | 0 | 0 | 0 | 0 | 0 | 0 | 0 |
| 1 | 0 | 0 | 0 | 0 | 245 | 0 | 0 | 0 | 0 | 0 | 0 | 0 | 0 | 0 | 0 | 0 | 0 | 0 | 0 | 0 | 0 | 0 | 0 | 0 |
| 2 | 0 | 809002 | 0 | 0 | 9574 | 0 | 0 | 0 | 0 | 0 | 0 | 0 | 0 | 0 | 0 | 0 | 0 | 0 | 0 | 0 | 21650 | 0 | 36754 | 0 |
| 3 | 0 | 0 | 0 | 0 | 0 | 0 | 0 | 0 | 0 | 0 | 0 | 0 | 0 | 0 | 0 | 0 | 87982 | 0 | 0 | 2589 | 0 | 0 | 0 | 0 |
| 7 | 0 | 0 | 0 | 0 | 0 | 0 | 0 | 0 | 0 | 0 | 0 | 0 | 0 | 0 | 0 | 0 | 0 | 0 | 0 | 0 | 0 | 0 | 0 | 0 |
| 8 | 0 | 0 | 0 | 0 | 0 | 0 | 0 | 0 | 0 | 0 | 0 | 0 | 0 | 0 | 0 | 0 | 0 | 0 | 0 | 0 | 0 | 0 | 0 | 0 |
| 9 | 0 | 0 | 0 | 0 | 0 | 0 | 0 | 0 | 0 | 0 | 0 | 0 | 0 | 0 | 0 | 0 | 0 | 0 | 0 | 0 | 0 | 0 | 0 | 0 |
| 10 | 0 | 0 | 0 | 0 | 0 | 0 | 0 | 0 | 0 | 0 | 0 | 0 | 0 | 0 | 0 | 0 | 39275 | 655 | 0 | 0 | 0 | 0 | 0 | 0 |
| 14 | 0 | 0 | 0 | 0 | 0 | 0 | 0 | 0 | 0 | 0 | 0 | 0 | 0 | 0 | 0 | 0 | 0 | 0 | 0 | 0 | 0 | 0 | 0 | 0 |
| 15 | 0 | 0 | 0 | 0 | 0 | 0 | 0 | 0 | 0 | 0 | 0 | 0 | 0 | 0 | 0 | 0 | 0 | 0 | 0 | 0 | 0 | 0 | 0 | 0 |
| 16 | 0 | 0 | 0 | 0 | 0 | 0 | 0 | 0 | 0 | 0 | 0 | 0 | 0 | 0 | 0 | 0 | 72908 | 0 | 0 | 0 | 0 | 0 | 0 | 0 |
| 17 | 0 | 0 | 0 | 0 | 0 | 0 | 0 | 0 | 0 | 0 | 0 | 0 | 0 | 0 | 0 | 0 | 0 | 0 | 0 | 0 | 0 | 0 | 0 | 0 |

#### Group 3[b]: JQ1 - JQ1 - vehicle

| Mouse | 25 | | 26 | | 27 | | 28 | | 29 | | 30 | | 31 | | 32 | | 33 | | 34 | | 35 | | 36 | |
|---|---|---|---|---|---|---|---|---|---|---|---|---|---|---|---|---|---|---|---|---|---|---|---|---|
| Day/eye[c] | L | R | L | R | L | R | L | R | L | R | L | R | L | R | L | R | L | R | L | R | L | R | L | R |
| 0 | 0 | 0 | 0 | 0 | 0 | 0 | 0 | 0 | 0 | 0 | 0 | 0 | 0 | 0 | 0 | 0 | 0 | 0 | 0 | 0 | 0 | 0 | 0 | 0 |
| 1 | 0 | 0 | 0 | 0 | 0 | 0 | 0 | 0 | 0 | 0 | 0 | 0 | 0 | 0 | 0 | 0 | 0 | 0 | 0 | 0 | 0 | 0 | 0 | 0 |
| 2 | 0 | 0 | 0 | 138118 | 0 | 0 | 0 | 0 | 0 | 0 | 0 | 0 | 0 | 277 | 107615 | 0 | 46499 | 0 | 0 | 0 | 0 | 0 | 194 | 0 |
| 3 | 0 | 0 | 0 | 13196 | 0 | 0 | 0 | 0 | 0 | 0 | 0 | 0 | 0 | 0 | 4415 | 0 | 18195 | 0 | 0 | 0 | 0 | 0 | 4447 | 408 |
| 7 | 0 | 0 | 0 | 0 | 0 | 0 | 0 | 0 | 0 | 0 | 0 | 0 | 0 | 0 | 0 | 0 | 0 | 0 | 0 | 0 | 0 | 0 | 0 | 0 |
| 8 | 0 | 0 | 0 | 0 | 0 | 0 | 0 | 0 | 0 | 0 | 0 | 0 | 0 | 0 | 0 | 0 | 0 | 0 | 0 | 0 | 0 | 0 | 0 | 0 |
| 9 | 0 | 0 | 0 | 0 | 0 | 0 | 0 | 0 | 0 | 0 | 297 | 0 | 0 | 0 | 0 | 1218 | 0 | 81523 | 4809 | 0 | 0 | 2640490 | 0 | 0 |
| 10 | 0 | 0 | 0 | 0 | 0 | 0 | 0 | 0 | 0 | 0 | 0 | 0 | 0 | 0 | 0 | 0 | 0 | 0 | 0 | 0 | 0 | 1430093 | 0 | 0 |
| 14 | 0 | 0 | 0 | 0 | 0 | 0 | 0 | 0 | 0 | 0 | 0 | 0 | 0 | 0 | 0 | 0 | 0 | 0 | 0 | 0 | 0 | 0 | 0 | 0 |
| 15 | 0 | 0 | 0 | 0 | 0 | 0 | 0 | 0 | 0 | 0 | 0 | 0 | 0 | 0 | 0 | 0 | 0 | 0 | 0 | 0 | 0 | 0 | 0 | 0 |
| 16 | 0 | 0 | 0 | 0 | 0 | 0 | 0 | 0 | 0 | 0 | 0 | 0 | 0 | 0 | 0 | 0 | 0 | 0 | 0 | 0 | 0 | 0 | 0 | 0 |
| 17 | 0 | 0 | 0 | 0 | 0 | 0 | 0 | 0 | 0 | 0 | 0 | 0 | 0 | 0 | 0 | 0 | 0 | 0 | 0 | 0 | 0 | 0 | 0 | 0 |

#### Group 4[b]: JQ1 - JQ1 - JQ1

| Mouse | 37 | | 38 | | 39 | | 40 | | 41 | | 42 | | 43 | | 44 | | 45 | | 46 | | 47 | | 48 | |
|---|---|---|---|---|---|---|---|---|---|---|---|---|---|---|---|---|---|---|---|---|---|---|---|---|
| Day/eye[c] | L | R | L | R | L | R | L | R | L | R | L | R | L | R | L | R | L | R | L | R | L | R | L | R |
| 0 | 0 | 0 | 0 | 0 | 0 | 0 | 0 | 0 | 0 | 0 | 0 | 0 | 0 | 0 | 0 | 0 | 0 | 0 | 0 | 0 | 0 | 0 | 0 | 0 |
| 1 | 0 | 0 | 0 | 0 | 0 | 0 | 0 | 0 | 0 | 0 | 0 | 0 | 0 | 0 | 0 | 0 | 0 | 0 | 0 | 0 | 0 | 0 | 0 | 0 |
| 2 | 0 | 0 | 0 | 0 | 0 | 0 | 0 | 0 | 0 | 0 | 332 | 0 | 1931180 | 987929 | 0 | 2530052 | 158478 | 0 | 0 | 0 | 0 | 0 | 0 | 0 |
| 3 | 0 | 0 | 0 | 0 | 0 | 0 | 0 | 0 | 0 | 0 | 0 | 0 | 14642 | 21781 | 0 | 3551 | 0 | 0 | 0 | 0 | 0 | 0 | 0 | 0 |
| 7 | 0 | 0 | 0 | 0 | 0 | 0 | 0 | 0 | 0 | 0 | 0 | 0 | 0 | 0 | 0 | 0 | 0 | 0 | 0 | 0 | 0 | 0 | 0 | 0 |
| 8 | 0 | 0 | 0 | 0 | 0 | 0 | 0 | 0 | 0 | 0 | 0 | 0 | 0 | 0 | 0 | 0 | 0 | 0 | 0 | 0 | 0 | 0 | 0 | 0 |
| 9 | 0 | 0 | 0 | 0 | 0 | 0 | 0 | 238238 | 472061 | 0 | 0 | 0 | 49403 | 0 | 194 | 38237 | 0 | 441625 | 0 | 157 | 4275968 | 0 | 0 | 5589 |
| 10 | 0 | 0 | 0 | 0 | 0 | 0 | 0 | 0 | 52991 | 0 | 0 | 0 | 0 | 0 | 0 | 120930 | 0 | 392154 | 0 | 0 | 35104 | 0 | 0 | 0 |
| 14 | 0 | 0 | 0 | 0 | 0 | 0 | 0 | 0 | 0 | 0 | 0 | 0 | 0 | 0 | 0 | 0 | 0 | 0 | 0 | 0 | 0 | 0 | 0 | 0 |
| 15 | 0 | 0 | 0 | 0 | 0 | 0 | 0 | 0 | 0 | 0 | 604 | 0 | 0 | 0 | 343 | 0 | 0 | 0 | 0 | 0 | 0 | 0 | 0 | 0 |
| 16 | 0 | 0 | 0 | 0 | 0 | 1541 | 0 | 0 | 0 | 0 | 0 | 0 | 0 | 0 | 1773082 | 0 | 0 | 0 | 0 | 0 | 0 | 0 | 0 | 9005 |
| 17 | 0 | 0 | 0 | 0 | 0 | 0 | 0 | 0 | 0 | 0 | 0 | 0 | 0 | 0 | 2442 | 0 | 0 | 0 | 0 | 0 | 0 | 0 | 0 | 208 |

[a] Virus titers are expressed in copies/ml.
[b] Mice were subjected to 3 weekly reactivation: mock reactivation (vehicle) for Group 1, JQ1 reactivation (50 mg/kg) for Group 4 or combinations of mock (vehicle) and JQ1 (50 mg/kg) reactivations for groups 2 and 3.
[c] At each time point, daily swabs were collected from day 0 to day 3 after each reactivation, from the left eye (L, blue) and right eye (R, orange) and analyzed separately.

**Supplemental Table 2.** Summary of Microscopic Findings in the Liver.

| | Naive[c,e] | HSV + AAV/MN[d,e,f] | AAV/MN only[c,e,f] |
|---|---|---|---|
| **No. animals examined** | **2** | **1** | **2** |
| **Hepatocellular phenotypic alteration** | (0)[a] | (1) | (2) |
| **Minimal** | - | - | - |
| **Mild** | - | 1[b] | - |
| **Marked** | - | - | 2 |
| **Inflammation mononuclear cell; Periportal and parenchymal** | (0) | (1) | (2) |
| **Minimal** | - | - | - |
| **Mild** | - | 1 | - |
| **Moderate** | - | - | 2 |
| **Hepatocellular necrosis, single cell** | (0) | (1) | (2) |
| **Minimal** | - | 1 | - |
| **Moderate** | - | - | 2 |
| **Hyperplasia; Kupffer and oval cell** | (0) | (1) | (2) |
| **Minimal** | - | - | - |
| **Moderate** | - | 1 | 2 |
| **Biliary cholestasis** | (0) | (1) | (2) |
| **Minimal** | - | - | - |
| **Mild** | - | - | 2 |

[a] Total number of animals with finding.
[b] Number of animals with the indicated severity for a specific finding.
[c] The liver was collected from naïve and AAV/MN only mice at 20 post AAV/MN administration.
[d] The liver was collected from HSV+AAV/MN mice at 33 post AAV/MN administration.
[e] All the mice were age-matched.
[f] AAV/MN was administered at a dose of 3.2 x $10^{12}$ vg AAV.

### **Supplemental Table 3.** Summary of Microscopic Findings in trigeminal ganglia.

| | CTRL no JQ1, n = 1[a,b] | | | CTRL 2xJQ1, n = 1[a,b] | | | AAV/MN[c] no JQ1, n = 3[a,b] | | | AAV/MN[c] 2xJQ1, n = 3[a,b] | | |
|---|---|---|---|---|---|---|---|---|---|---|---|---|
| **1- Trigeminal ganglia** | Score | Mean score[d] | Mean severity[e] | Score | Mean score[d] | Mean severity[e] | Score | Mean score[d] | Mean severity[e] | Score | Mean score[d] | Mean severity[e] |
| Inflammation, L/P/E | 1 | 1 | 1 | 1 | 1 | 1 | 3 1 1 | 1.6 | 1.6 | 2 1 2 | 1.6 | 1.6 |
| Neuronal degeneration | 0 | 0 | 0 | 0 | 0 | 0 | 2 0 0 | 0.7 | 2 | 1 0 1 | 0.7 | 2 |
| Neuronal necrosis | 0 | 0 | 0 | 0 | 0 | 0 | 1 0 1 | 0.7 | 1 | 1 0 1 | 0.7 | 2 |
| Perivascular infiltrates | 0 | 0 | 0 | 0 | 0 | 0 | 1 0 0 | 0.3 | 1 | 0 0 0 | 0 | 0 |
| **Sum - Scores:** | **1** | **1** | **1** | **1** | **1** | **1** | **7 1 2** | **3.3** | **5.6** | **4 1 4** | **3** | **5.6** |
| **2- Associated nerve fiber** | Score | Mean score[d] | Mean severity[e] | Score | Mean score[d] | Mean severity[e] | Score | Mean score[d] | Mean severity[e] | Score | Mean score[d] | Mean severity[e] |
| Dilated myelin sheath | 2 | 2 | 2 | 2 | 2 | 2 | 3 3 3 | 3 | 3 | 2 1 2 | 1.7 | 1.7 |
| Digestion chamber | 0 | 0 | 0 | 1 | 1 | 1 | 2 2 0 | 1.3 | 2 | 1 0 1 | 0.7 | 1 |
| Perivacular infiltrates | 0 | 0 | 0 | 0 | 0 | 0 | 2 1 2 | 1.7 | 1.7 | 1 0 1 | 0.7 | 1 |
| Inflammation, L/P/E | 1 | 1 | 1 | 1 | 1 | 1 | 1 1 3 | 1.7 | 1.7 | 0 0 0 | 0 | 0 |
| **Sum - Scores:** | **3** | **3** | **3** | **4** | **4** | **4** | **8 7 8** | **7.7** | **8.4** | **4 1 4** | **3.1** | **3.7** |

[a] Number of TG per slide
[b] Each slide had 3 sections from one TG
[c] AAV dose administered was 1.8 x$10^{12}$ vg/mouse
[d] Mean score = Sum of scores from TG in the group/number of TG in the group
[e] Mean severity = Sum of scores from TG in the group/number of TG with a score > 0 in the group
